# Interplay of the ribosome A and CAR sites

**DOI:** 10.64898/2026.04.07.714784

**Authors:** Mitsu Raval, Yancheng Zhou, Miki Lynch, Daniel Krizanc, Kelly M. Thayer, Michael P. Weir

## Abstract

Protein translation is a highly regulated process influenced by multiple factors at the initiation, elongation, and termination stages. One notable regulatory element of the ribosome is the CAR interaction surface, a three-residue motif in the structure of the ribosome composed of C1274 and A1427 of *S. cerevisiae* 18S rRNA (corresponding to C1054 and A1196 in *E. coli* 16S rRNA) and R146 of ribosomal protein Rps3. CAR is highly conserved and positioned adjacent to the amino-acyl (A site) decoding center. It establishes hydrogen bonds with the +1 codon next in line to enter the ribosome A site, acting as an extension of the tRNA anticodon and forming base-stacking interactions with nucleotide 34 of the tRNA. However, despite CAR’s enzymatically strategic positioning within the ribosome, its functional relationship with the A site remains poorly characterized. Using molecular dynamics (MD) simulations, we examined the interplay between the A site and CAR site, revealing sequence-dependent modulation of H-bonding and π-stacking interactions within and between the two sites. These findings highlight the interplay between the A site and CAR site, suggesting a structural and functional connection between these two regions of the ribosome that may contribute to mRNA sequence-specific tuning of translation elongation.

**GRAPHICAL ABSTRACT:** 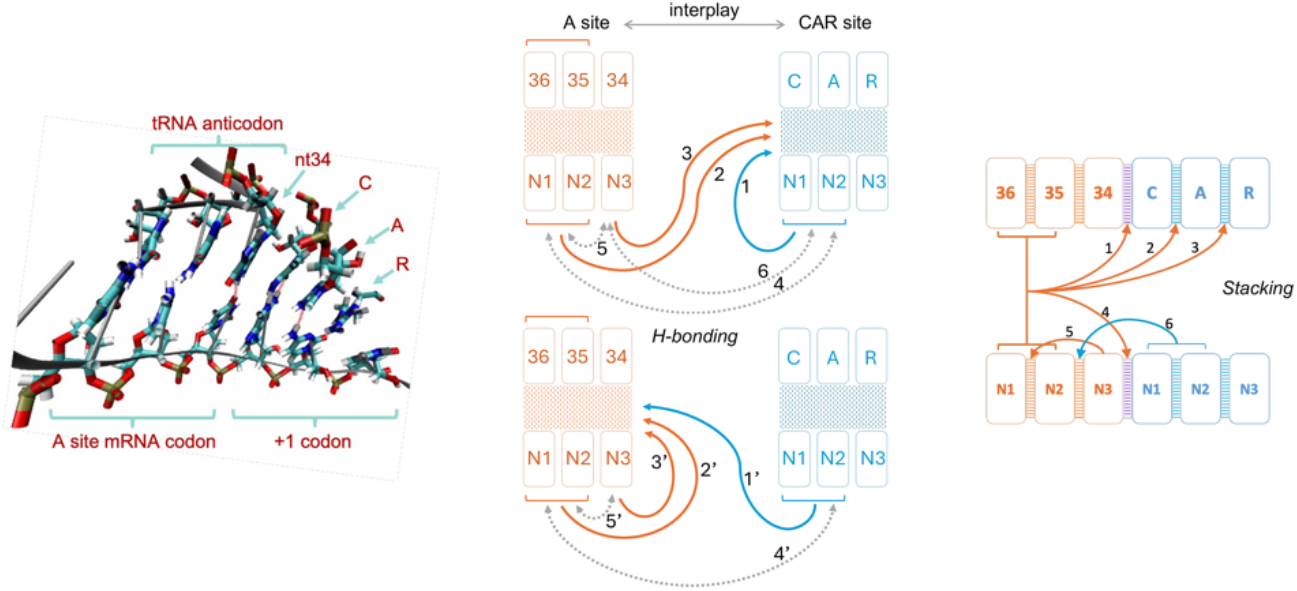

## INTRODUCTION

Protein translation is a tightly regulated process orchestrated by the ribosome. Translation has typically been described as the decoding of individual mRNA codons to synthesize a growing polypeptide chain of amino acids(1–3). Various motifs within the structure of the ribosome are known to regulate the codon recognition efficacy and rate of mRNA threading in translation(4–12). While classical models promote a codon-by-codon mechanism in decoding of translation, the hypothesis that adjacent codon context provides cis-regulatory influences in translation has gained support in recent years(13–15). The CAR interaction surface is a regulatory motif in the structure of ribosome that has been implicated in integrating codon adjacency effects in translation(16, 17) (Fig. 1).

**Figure 1.**
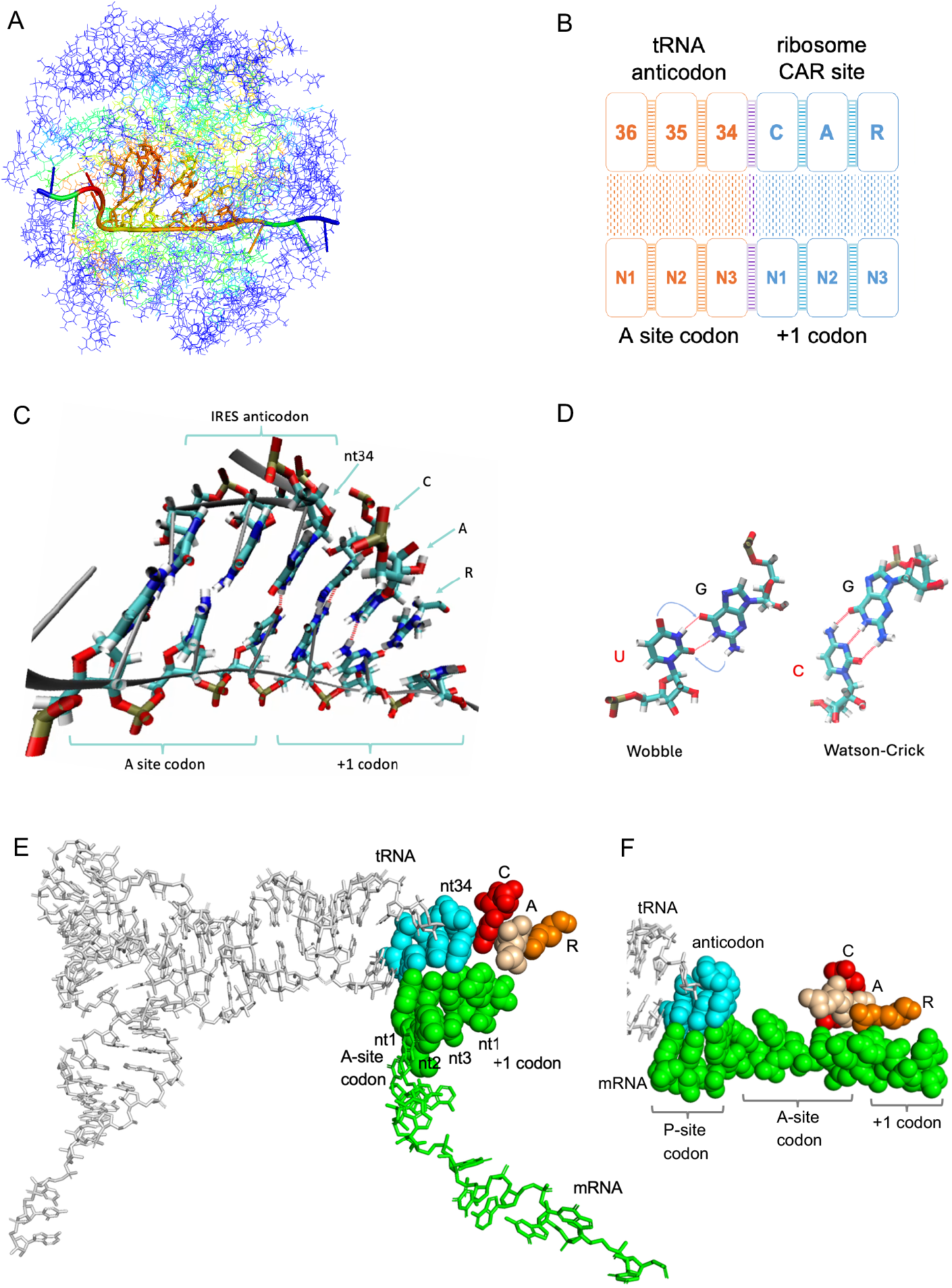
CAR forms sequence specific interactions with the mRNA +1 codon potentially regulating ribosome translation. (**A**) The yeast ribosome translocation stage II structure (PDB ID 5JUP),(18) in which a viral IRES mimics the mRNA and tRNAs, was used to create a 494 residue decoding center neighborhood subsystem centered around the C of CAR. The outer residues colored in blue were restrained to maintain the translocation stage II structure during molecular dynamics (MD) simulations to observe the behavior of the central CAR and A sites.(16) Residues are colored based on their root mean square fluctuation (RMSF) values. The outer “onion shell” residues in dark blue do not fluctuate while the inner core residues move freely with the most fluctuating residues in red. (**B**) Schematic representation of the A site and the CAR site. Hydrogen bonds (H-bonds) are represented as a stippled pattern and stacking as ladder pattern between neighboring residues. (**C**) VMD Licorice representation of an MD simulation frame that includes the A site, CAR, and mRNA +1 codon. CAR behaves like an extension of the A site tRNA anticodon. CAR is anchored to the anticodon through stacking between tRNA nucleotide 34 and C1054 of CAR. CAR H-bonds (red dashes) with the mRNA +1 codon(22). (**D**) tRNA G34 can base pair with U or C via wobble or Watson-Crick base pairing respectively^3^. (**E**) Early translocation structure (PDB ID 7RR5) with mRNA and tRNAs instead of an IRES, showing CAR (C, red; A, wheat, R, orange) as an extension of the tRNA anticodon(20). The A-site codon and first nucleotide of the +1 codon are resolved in the structure. CAR is positioned close to the anticodon in early translocation intermediates and less close in later intermediates (Fig. S1). (**F**) Preinitiation structure (PDB ID 6FYX(21)) with Met-tRNA accommodated in the P site and CAR abutting A-site nt 3, +1 codon nt 1 and nt 2; also see Fig. S2. Genetic studies suggest that R of CAR may participate in selection of translation initiation sites(35).

CAR was first discovered in cryo-EM structures of early translocation stages in yeast ribosomes formed with Taura syndrome virus IRES (internal ribosome entry site) and translocase cEF2*GTP bound with sordarin which closely resemble canonical translation (Fig 1C) (16, 18). CAR has since been observed in translocation intermediates with mRNA and tRNAs instead of an IRES (Fig 1E, Fig. S1(19, 20)) as well as preinitiation structures (Fig 1F, Fig. S2(21)). In this study, we use an IRES-containing translocation stage structure to investigate CAR function. CAR is comprised of the two nucleotides C1274 and A1427 of 18S rRNA (which we refer to as C1054 and A1196, the corresponding residues in *E. coli* 16S rRNA), and an amino acid R146 of the Rps3 protein in the *S. cerevisiae* ribosome. C1054 and A1196 are well conserved across prokaryotes and eukaryotes and R146 is conserved across eukaryotes. C and A belong to two separate loops of 18S rRNA helix 34, whereas R is contributed by a finger extension of the Rps3 protein. The assembly and folding of the ribosome support the structural integrity of CAR that emerges from the stacking of the C, A, and R residues through pi-pi (C:A) and pi-cation (A:R) interactions, similar to stacking in the adjacent tRNA anticodon (Fig. 1B, Fig. 1C). Indeed, CAR behaves as an extension of the anticodon with C of CAR stacking with nucleotide 34 of the tRNA, the nucleotide that base pairs with the wobble nucleotide of the A-site codon (Fig 1C, E). The stacking is pronounced in the early stages of translocation and less so at later stages when the CAR surface moves away from the anticodon(16) (Fig. S1).

This positioning of CAR adjacent to the A-site anticodon ideally situates CAR so that its residues can H-bond to the stacked residues of the +1 codon—the codon 3’ adjacent to the A-site codon—using the Watson-Crick edge of the C base (of CAR), the Hoogsteen edge of the A base, and an edge of the planar guanidinium group of R (Fig. 1C)(16, 22). We define the interacting CAR and +1 codon residues as the ribosome’s “CAR site.” In the adjacent A site, nucleotide 34 of the tRNA anticodon has the structural flexibility of forming either wobble or Watson-Crick base pairing depending on the identity of the wobble-position nucleotide in the A-site mRNA codon, potentially influencing the relationship between the A and CAR sites (Fig. 1D).

The regulatory potential of the CAR site resides in its enzymatically strategic positioning and sequence specificity. The levels of H-bonding at the CAR site are higher when the +1 codon is GNN (where N is A, C, G or U), particularly when 3’ of an A-site NNU codon (denoted by the codon context aNNU_+1GNN)(17, 22). Correspondingly, ribosome profiling analysis has revealed slower mRNA threading, indicated by higher ribosome densities, in aNNU_+1GNN mRNA sequence contexts(17). This suggests a model where CAR’s H-bonding with the mRNA acts as a “braking system” that slows down the rate of mRNA threading in protein synthesis based on the mRNA sequence context of adjacent A- and CAR-site codons, and in particular, the identities of the A-site wobble nucleotide, and the first nucleotide of the +1 codon(17).

This paper explores further the potential role of the CAR surface in integrating the cis-regulatory codon-adjacency signals between the CAR and A sites. Previous mutation and suppressor studies have implicated C of CAR in codon recognition and frame maintenance functions that have been attributed solely to effects at the A site(23–28). Moreover, various A-site targeting molecules such as the bacterial toxin Hig B, ribosome associated enzymes, antibiotics tetracycline, tigecycline and streptothricin F, include C and/or A of CAR in their binding pockets(29–34). R of CAR has also been implicated in A-site decoding and translational initiation in mechanisms exploited by the SARS CoV2 virus(23). These studies have revealed targeting of A-site functionality through interactions that include residues of CAR, potentially exploiting structural and mechanistic communication between the CAR and A sites. However, this communication remains unexplored, and a better understanding of the interplay between the evolutionarily conserved A and CAR sites could open novel perspectives for investigating adjacent codon cis-regulation, structure informed therapeutic intervention, and evolutionary codon biases of mRNA open reading frame (ORF) sequences.

Using molecular dynamics (MD) simulations, we explore the mechanistic structural interplay between the A site and CAR site through quantitative analysis of H-bonding and stacking interactions within and between both sites. Our results suggest that the CAR interface contributes as a central player in the sequence-dependent structural interplay between codons at the adjacent sites during protein translation.

## METHODS

### Molecular dynamics simulation system setup

We employed a 494-residue ribosome subsystem centered around C of CAR, encompassing the decoding center and the CAR interaction surface^6^. Coordinates were derived from the translocation stage II (PDB: 5JUP) structure of the yeast cryo-EM analysis which had originally been used to resolve five discrete structural stages of ribosome translocation through computational integration of 1.1M two-dimensional images of ribosomes(18). The subsystem contained 183 nucleotides and 311 amino acids, designed to include the A site codon-anticodon, the CAR interaction residues, and the adjacent +1 codon. The structure also incorporates a minimal mRNA-tRNA complex, modeled as an internal ribosome entry site (IRES) stem-loop from Taura syndrome virus that reproduces the codon–anticodon interaction (IRES pseudoknot I, PKI) without requiring the full tRNA body. An “onion shell” restraint was used to stabilize peripheral residues during dynamics while leaving the core—including the A-site, CAR surface, and +1 codon—flexible. Specifically, a 20 kcal/mol·Å^2^ positional restraint was applied to the 173 outermost residues, with a buffer region of at least one restrained residue separating the flexible core from the solvent. Mutations to the A-site and +1 codon triplets, as well as the anticodon, were introduced using AMBER’s tLEaP utility, maintaining the maximum number of experimental atomic coordinates and regenerating missing atoms to preserve local geometry^32^.

### Molecular dynamics simulations

Simulations were performed as described previously(36) using the AMBER ff14SB force field for proteins and ff99bsc0χOL3 for RNA, in conjunction with the TIP3P water model and potassium counterions to neutralize system charge(37–39). A 12 Å solvent box with periodic boundary conditions was used. Following energy minimization and heating, all systems were equilibrated at 300 K before running production dynamics. Each mRNA:tRNA combination was simulated using 30 replicate trajectories: 20 of 60 ns and 10 of 100 ns, totaling 2.2 μs of simulation per condition. Analysis was restricted to frames from the final 40 or 80 ns of 60 or 100 ns trajectories respectively, based on root mean square deviation (RMSD) stabilization assessments using cpptraj(40) which typically showed average deviations of backbone atoms from the starting structure of 2Å or less.

### Structural feature quantification

Hydrogen bonding was quantified using cpptraj’s hbond function with a default donor–acceptor distance cutoff of 3.5 Å and an angle cutoff of 135°, applied to selected atom pairs between nucleotides of interest (A-site, +1 codon, tRNA anticodon, CAR). For stacking analysis, center-of-geometry (COG) distances were calculated between relevant nucleobases or, in the case of the A:R stack, between the purine base and the guanidinium group of R146(17, 36, 41). Purine base centers were selected as the minimum COG from either the imidazole or pyrimidine ring. Ten stack-pairs were selected for analysis, spanning internal codon, anticodon, CAR-site, and cross-interface interactions. COG values served as structural proxies for π–π and π–cation stacking strength, with lower values indicating more stable stacking.

### Data aggregation and statistical analysis

Stacking and H-bonding metrics were compiled across all replicates using custom Python and R scripts enhanced by generative AI (ChatGPT version 4o). For each feature, mean values were calculated per structure and per condition (e.g., wobble vs. Watson–Crick, +1GCU vs. +1CGU). To perform multivariate analysis multifactorial ANOVA models were used on data from 30 replicates to assess statistical significance of codon identity effects(42). The p-values were adjusted for multiple comparisons using the Bonferroni correction to reduce the likelihood of false-positive results arising from repeated hypothesis testing(43). To visualize higher-order organization, correlation matrices and hierarchical clustering were performed using Pearson’s r between H-bonding or stacking vectors across structures. Violin plots, dot plots, and annotated schematic diagrams were produced to illustrate the geometry and crosstalk of interactions spanning the codon– anticodon–CAR–+1 region.

### Clustering analysis of interaction profiles

For interaction-level UMAP (Uniform Manifold Approximation and Projection; Fig. S9 and 4A) and PCA (Principal Component Analysis; Fig. S10), each of the twenty-six contacts (a–z) was represented by a 720-element vector of mean interaction values collected from all post-equilibration runs (24 structures × 30 replicates = 720 total simulations; MD runs for Figs 2B and 2D). H-bonding and stacking measures were scaled by variance to allow cross-comparisons. We then computed Pearson’s r between every pair of these 720-dimensional interaction vectors and converted each correlation into a distance via “1 – r,” producing a 26 × 26 correlation-distance matrix. UMAP was applied directly to this distance matrix with n neighbors = 15, min dist = 0.1, and metric = “precomputed”. The resulting two-dimensional embedding arranges interactions (a–z) according to their covariation patterns across all post-equilibration runs, enabling visual identification of which contacts behave similarly over the entire ensemble.

**Figure 2.**
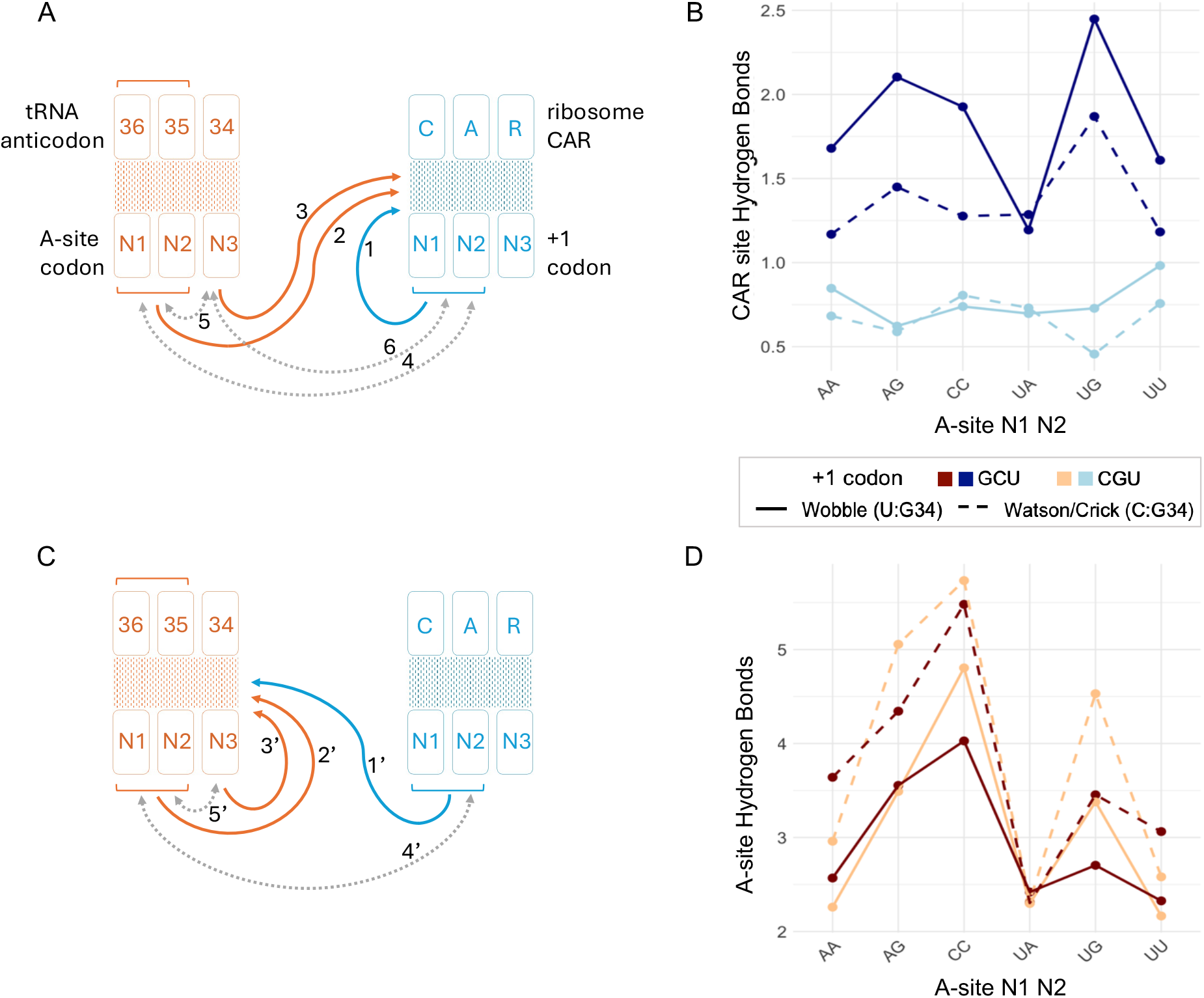
Interplay between the A site and CAR site revealed through ANOVA analysis of H-bonding. (**A, C**) Schematic representation of the decoding center highlighting how local codon context influences H-bonding at the CAR site (A) and the A site (C). Solid arrows (1–3, 1′–3′) represent the three main effects: the impact of +1 codon identity (1, 1′), A-site codon N1N2 identity (2, 2′), and A-site wobble decoding geometry at position N3 (3, 3′) on CAR-site (A) and A-site (C) H-bonding. Dotted arrows (4–6, 4’-5′) depict two-way interactions between variables: A-site codon and +1 codon modulate each other’s effects (4), wobble geometry and A-site N1N2 modulate each other’s effects (5), and wobble geometry and +1 codon modulate each other’s effects (6) on CAR-site H-bonding; with analogous interactions (4’, 5′) tuning A-site H-bonding. All arrows have *p* < 0.001 except arrow 1′ (*p* < 0.01) and arrow 5 (*p* < 0.05). (**B, D**) Graphical representation of mean H-bond counts across A-site codons (x-axis), comparing GCU and CGU +1 codon contexts (color: dark blue vs. light blue in B, dark red vs. light orange in D) and decoding geometries (solid = wobble G:U; dashed = Watson–Crick G:C). Upper plots (B) show CAR-site H-bonding; lower plots (D) show A-site H-bonding. As described previously, for each structure we quantified the strength of H-bonding at each site by calculating the mean frequency of H-bonds formed by mRNA nucleotides with the three nearest residues on their complementary surface in the trajectory frames of MD simulations (30 MD replicates for each structure)(22). Positions 36 and 35 of the tRNA anticodon were substituted to maintain Watson–Crick complementarity with A-site codon positions N1 and N2, ensuring consistent pairing across all constructs. H-bonding data of CAR : +1 codon interactions for A-site N1N2 codons aAG, aCC, aUG, and aUU shown in (B) were previously reported in Sun et al. (17).

For structure-level UMAP (Fig. S9), each of the 720 post-equilibration simulations was instead represented by its own 26-element interaction fingerprint (one entry per a–z contact summarized in Fig. 4A). We computed Pearson’s r between every pair of these 720 vectors and again converted to distance via “1 – r”, yielding a 720 × 720 correlation-distance matrix. UMAP was then run on that full matrix using the same hyperparameters, producing a two-dimensional embedding in which each point corresponds to one simulation replicate’s overall interaction profile.

### Covariation analysis between interactions

To resolve interaction coupling at a finer resolution, we performed a frame-resolved correlation analysis using custom Python scripts. For each MD replicate, H-bond counts and stacking geometries were extracted for every trajectory frame after equilibration. To exclude non-equilibrated dynamics, the first 2000 frames of each trajectory were removed prior to analysis.

Stacking interactions were converted from continuous center-of-geometry (COG) distances into binary stacking events using a cutoff of 4.5 Å, where distances ≤ 4.5 Å were classified as stacked and distances > 4.5 Å as unstacked, consistent with prior structural definitions of stable π–π and π–cation stacking(41). H-bond presence was also treated as a binary variable per frame using standard cpptraj distance (3.5 Å) and angle (135°) criteria.

Because both variables were binary at the frame level, φ (phi) correlation coefficients were computed to quantify association between stacking and H-bonding events across frames. φ correlation is mathematically equivalent to Pearson’s r for binary data and provides a normalized measure of association bounded between −1 and +1. This approach enables detection of direct coupling between discrete interaction events, rather than correlations driven by averaged magnitudes.

In parallel, we constructed a 26 × 26 covariation matrix by calculating across all post-equilibration runs for every pair of interactions (a–z). This matrix was visualized as a heatmap, highlighting all cross-site covariations, and informing which contacts co-vary most strongly.

## RESULTS AND DISCUSSION

Translation of proteins is a result of a highly coordinated and sensitive structural communication network formed by the ribosome, tRNAs, mRNA and other enzymes. Codon adjacency effects might influence this network to further tune the regulation of translation(13–15). To investigate how local mRNA sequence context can modulate residue-residue interactions in the decoding center neighborhood during translation, we quantified hydrogen bonding (H-bonding) interactions at both the CAR site and the A site (Fig. 1B, C) using MD simulations of a viral mRNA/yeast ribosome subsystem performed under systematically varied mRNA configurations. H-bonds were quantified by averaging H-bond occurrences across the MD trajectory frames. At the CAR site, we measured H-bonds formed between the ribosomal CAR surface and the +1 codon; at the A site, we quantified H-bonding between A-site codons and the anticodon of a G34-containing tRNA.

In both cases, we varied three factors: (i) the dinucleotide identity at positions 1 and 2 of the A-site codon, (ii) the A-site wobble (third) nucleotide identity and geometry (see below), and (iii) the identity of the CAR-site +1 codon (+1GCU or +1CGU). In all decoding configurations, tRNA anticodon positions 36 and 35 (Fig. 1B) were substituted to complement A-site nucleotides 1 and 2 (N1 and N2), while position 34 was always guanosine to enable either wobble (U:G34) or Watson–Crick (C:G34) geometry base pairing with the third nucleotide of the A-site codon (Fig. 1D). We focused this study on NNU and NNC codons that utilize G34 tRNAs (12 codons in yeast; Fig. 2B) because previous preliminary analysis suggested that these codon/anticodon and +1 codon combinations are particularly influential in differentially affecting ribosome profile and MD behaviors.

30 independently initialized replicates were performed for each version of our MD system. However, because H-bond counts were not normally distributed across the replicates (Fig. S3), we used aligned rank transform (ART) ANOVA to assess main effects and interactions(44). Residuals from classical ANOVA failed the Shapiro– Wilk tests for both datasets, justifying the use of a non-parametric approach^35^. This experimental design is outlined in Figure 2A and the statistical findings summarized in Tables S1, S2 and S3.

### CAR as a molecular decoder of codon context

Based on the Bonferroni corrected ART-ANOVA analysis shown in Figure 2, CAR-site H-bonding was shaped by each of the three main variables: A-site N1 N2, wobble nucleotide identity and the CAR site +1 codon sequence. A-site mRNA codon nucleotides 1 and 2 (N1 N2) identity exerted a strong effect (p < 2 × 10^−16^), with certain A-site codons supporting greater CAR interaction with the +1 codon (arrow 2, Fig. 2A; Fig. 2B). For example, aUGU_+1GCU (red arrowhead, Fig. S4B) formed significantly more CAR H-bonds than aUAU_+1GCU (blue arrowhead, Fig. S4B). Consistent with prior observations(17), wobble decoding geometry at the A-site wobble position (Fig 1D) further enhanced CAR interactions (arrow 3, Fig. 2A); e.g., aAGU_+1GCU (red arrowhead, Fig. S4C) exhibited stronger CAR H-bonding than aAGC_+1GCU (blue arrowhead, Fig. S4C), reflecting how A-site decoding geometry modulates downstream +1 codon engagement (Fig. 2B, Fig. S4C). Consistent with previous observations(22), the identity of the +1 codon also played a major role (p = 0.0113) where +1GCU supported stronger CAR interactions than +1CGU (arrow 1, Fig. 2A), particularly when the A-site wobble nucleotide partakes in wobble base pairing (G:U); e.g., aUGU_+1GCU (red arrowhead, Fig. S4D) showed higher H-bonding than aUGU_+1CGU (blue arrowhead). Collectively, these results indicate that A-site codon composition, including both codon identity and decoding geometry, exerts a significant influence on CAR-mediated H-bonding to the +1 codon, particularly in +1GCU contexts. This supports the hypothesis that the A site modulates translational rates in part by tuning the strength of CAR’s engagement with the downstream +1 codon.

In addition to these main effects, two-way interactions of variables (grey two-way arrows, Fig. 2A) revealed that CAR acts as an integrative surface that fine-tunes how codon adjacency influences manifest. Interactions between A-site codons N1 N2 and +1 codons (arrow 4), A-site codons N1 N2 and N3 wobble pairing (arrow 5), and +1 codons and N3 wobble pairing (arrow 6), modulated the main effects shown by arrows 1, 2, and 3 (Fig. 2A).

For example, the difference in CAR H-bonding between +1GCU and +1CGU (arrow 1) was amplified (arrow 4) when the A-site codon was UGU (red arrowheads, Fig. S4E) compared to UAU (blue arrowheads), highlighting how A-site codon identity tunes the +1 codon effect. Conversely, the +1 codon context (arrow 4) gated the main effect of arrow 2 (i.e., the influence of A-site sequence on CAR H-bonding) as seen in comparisons aAGU_+1GCU > aAAU_+1GCU (red arrowheads, Fig. S4F) versus aAGU_+1CGU ≈ aAAU_+1CGU (blue arrowheads).

Similarly, wobble pairing (arrow 5) modulated arrow 2 (the effect of A-site codon identity on CAR engagement), as illustrated by comparisons aCCU_+1GCU vs. aUAU_+1GCU (red arrowheads, Fig. S4G) versus aCCC_+1GCU vs. aUAC_+1GCU (blue arrowheads). In these cases, wobble decoding amplified A-site codon differences, suggesting that decoding geometry can selectively enhance sequence-specific CAR sensitivity. Conversely, wobble-induced gains in CAR H-bonding (arrow 3) were more pronounced for some A-site codons (e.g., CCU; red arrowheads, Fig. S4H) than others (e.g., UAU; blue arrowheads), indicating that the extent to which wobble pairing affects CAR engagement is contingent on the upstream A-site N1 N2 codon sequence (arrow 5).

Moreover, the impact of wobble pairing on CAR H-bonding itself (arrow 3) was dependent on the +1 codon identity (arrow 6). For example, the wobble-dependent boost in CAR bonding was more pronounced when +1GCU was present (aCCU_+1GCU vs. aCCC_+1GCU; red arrowheads, Fig. S4I) compared to +1CGU (aCCU_+1CGU vs. aCCC_+1CGU; blue arrowheads). Conversely, +1 codon effects (arrow 1) were also shaped by the decoding geometry (arrow 6) where comparisons such as aCCU_+1GCU vs. aCCU_+1CGU (red arrowheads, Fig. S4J) versus aCCC_+1GCU vs. aCCC_+1CGU (blue arrowheads) revealed that the +1 codon’s influence on CAR engagement was heightened under wobble decoding.

Together, these results reinforce CAR’s likely role as a context-sensitive molecular decoder of codon adjacency, that integrates A-site codon identity, decoding geometry, and downstream codon context to fine-tune its own H-bonding engagement with the +1 codon.

### The CAR site interactions modulate A-site H-Bonding through contextual cues

Having established that CAR’s H-bonding interactions are influenced by both +1 codon and A-site sequences, we next investigated whether this relationship is reciprocal. Specifically, we wanted to determine if the CAR site (CAR : +1 codon region) could in turn, affect H-bonding at the A site based on codon adjacency context. We analyzed how A-site H-bonding was affected by the A-site mRNA codon and the downstream +1 codon sequence variables. Three main effects emerged (arrows 1′, 2′, and 3′, Fig. 2C, 2D): the identity of the +1 codon, the identity of the A-site N1 N2 bases, and the wobble decoding geometry at N3.

The CAR site’s influence reached upstream, altering interactions not just at its own site, but at the A-site decoding center itself. The identity of the +1 codon significantly modulated A-site H-bonding (arrow 1’, Fig. 2C): for instance, aUGC_+1CGU (red arrowhead, Fig. S5B) supported more H-bonds than aUGC_+1GCU (blue arrowhead), despite identical tRNA base pairing potential at the A site (Fig. 2D). This demonstrates an upstream directed communication channel, wherein the CAR site transmits sequence-dependent signals to reshape A-site geometry—a form of tuning of the CAR and tRNA interactions based on the local mRNA sequence, integrating adjacent codon context into the mechanics of translation.

A bidirectional crosstalk model was further supported by two-way interactions between variables: the effect of the +1 codon (arrow 1’) depended on A-site identity (arrow 4’ Fig. 2C). Comparing aAGC and aAAC codons, the difference between +1CGU and +1GCU was larger for aAGC (red arrowheads, Fig. S5C) than for aAAC (blue arrowheads), indicating that certain A-site codons are more sensitive to the +1 codon’s regulatory effect than others. Conversely, the main effect of A-site codon identity (arrow 2′) also varied with the +1 codon context (arrow 4’): the difference between aAGC and aAAC was greater when paired with +1CGU (red arrowheads, Fig. S5D) than with +1GCU (blue arrowheads), showing that the downstream +1 codon sequence tunes the ability of the A site to engage the tRNA anticodon.

As expected, A-site codon identity at positions N1 and N2 strongly influenced A-site H-bonding (arrow 2′, Fig. 2C), consistent with canonical base-pairing energetics: codons forming G:C base pairs, such as aCCC (red arrowhead, Fig. S5E), supported more H-bonds than codons forming A:U base pairs, such as aUAC (blue arrowhead). Wobble base pairing at the third codon position (N3) also modulated A-site H-bonding (arrow 3′, Fig. 2C). Watson–Crick G:C pairs between the wobble A-site N3 nucleotide and the nucleotide 34 of the tRNA anticodon (e.g., aCCC_+1GCU, red arrowhead, Fig. S5F) generally supported higher H-bond counts than G:U wobble pairs (e.g., aCCU_+1GCU, blue arrowhead, Fig. S5F). The agreement of both N1 N2 and N3 decoding trends with established base-pairing principles serves as an internal validation of our H-bonding analysis framework.

Our results so far reveal a structural interplay between the ribosome’s A site and its downstream adjacent CAR site. Specifically, we observed that the mRNA codon sequence at the A site affects the H-bonding functionality at the CAR site (Fig. 2A). Similarly, the mRNA +1 codon sequence at the CAR site also affects the H-bonding between the A-site codon and its anticodon (Fig. 2C). However, we were curious if the H-bonding effects at these sites were simply dependent on the mRNA sequence nucleotide identities alone irrespective of CAR’s influence or if CAR’s engagement with the mRNA—the hypothesized CAR braking system—altered this mechanistic structural interplay. We analyzed the correlation between H-bonding at the CAR site and the A site to explore these possibilities.

We first examined the overall correlation between H-bonds at the two sites (Fig. S6A). A very weak but statistically significant anticorrelation was observed between the two sites. Specifically, higher A-site H-bonding tended to coincide with weaker CAR engagement.

To further dissect how mRNA sequence context shapes this correlation of H-bonds between both sites, we stratified the analysis by A-site codon identity (Fig. S6B). We computed the Pearson correlation coefficient between A-site and CAR-site H-bonds for different A-site codons separately with CGU and GCU +1 codons. Across most A-site codons, weak negative correlations were observed. However, the strength and direction of this relationship varied considerably depending on the adjacent A-site and +1 codon sequences.

Notably, A-site UU codons (UUU or UUC) in conjugation with +1GCU exhibited the strongest negative correlation under +1GCU conditions (r = -0.428, p < 7.87 × 10^−3^ after Bonferroni correction, Table S4), revealing anticorrelation between A- and CAR-site H-bonding when a U-rich A-site codon precedes a GCU +1 codon. In other A-site contexts (e.g., AA or CC), +1GCU tended toward negative r values but did not reach significance with multiple testing. This stratified analysis emphasizes that the coupling of behaviors at the adjacent A and CAR sites is highly sequence dependent.

Taken together, the weak correlations observed suggest a potential H-bond–based interplay between the two sites that may be subtle and context-specific. However, because we calculated correlations on mean A-site and mean CAR-site H-bond counts, any bond-specific positive and negative associations are obscured. In practice, this aggregation may cancel out individual trends resulting in an overall Pearson r that underestimates finer-scale interactions. Moreover, this H-bond-only model omits π-stacking effects, which are known to contribute substantially to ribosome structural dynamics(41).

### CAR-mediated neighborhood interplay interprets and modulates adjacent codon neighborhood architecture

The above analysis raised the idea that the CAR surface plays a role in mediating interactions between the adjacent A- and CAR-site codons. However, it remains possible that the observed interplay effects between the A-site and CAR-site are not actively mediated through CAR itself but are instead largely transmitted through stacking of the mRNA nucleotides across these two sites. To test this hypothesis, we examined π-stacking interactions of adjacent residues using distances between their center-of-geometry (COG of base rings) as a structural proxy for stacking strength, with lower values corresponding to tighter, more stable stacking (< 4.5Å)(36, 41). We reported the mean COG values calculated for each simulation. Stacking interactions were measured between nucleotide bases, except for the A-to-R interaction of CAR, which reflected π–cation stacking between adenine and the guanidinium group of arginine(36).

The results, shown in Figure 3, revealed widespread modulation of stacking geometry by local sequence context. We observed statistically significant main effects and interactions across numerous stack pairs (Table S5). To accommodate the non-normal data distributions shown in Fig. S7, we used the nonparametric ART-ANOVA approach. We adjusted for multiple comparisons using Bonferroni correction. To interpret these stacks, we introduce a hierarchy of zero- and non-zero-order effects, where zero-order effects are defined as a nucleotide modulating its own stacking with a neighbor (e.g., A-site N2 effects on the N2:N3 stacking; Fig. 3A) and non-zero-order effects reflect modulation one or more nucleotides away (e.g., A-site N1 N2 affects stacking within the CAR motif; arrows 2 and 3, Fig. 3A). While zero-order effects aligned with expectations and served to validate our model, it was the non-zero-order effects—illustrated by the arrows in Figure 3A—that uncovered structural coordination across the adjacent codon landscape.

**Figure 3.**
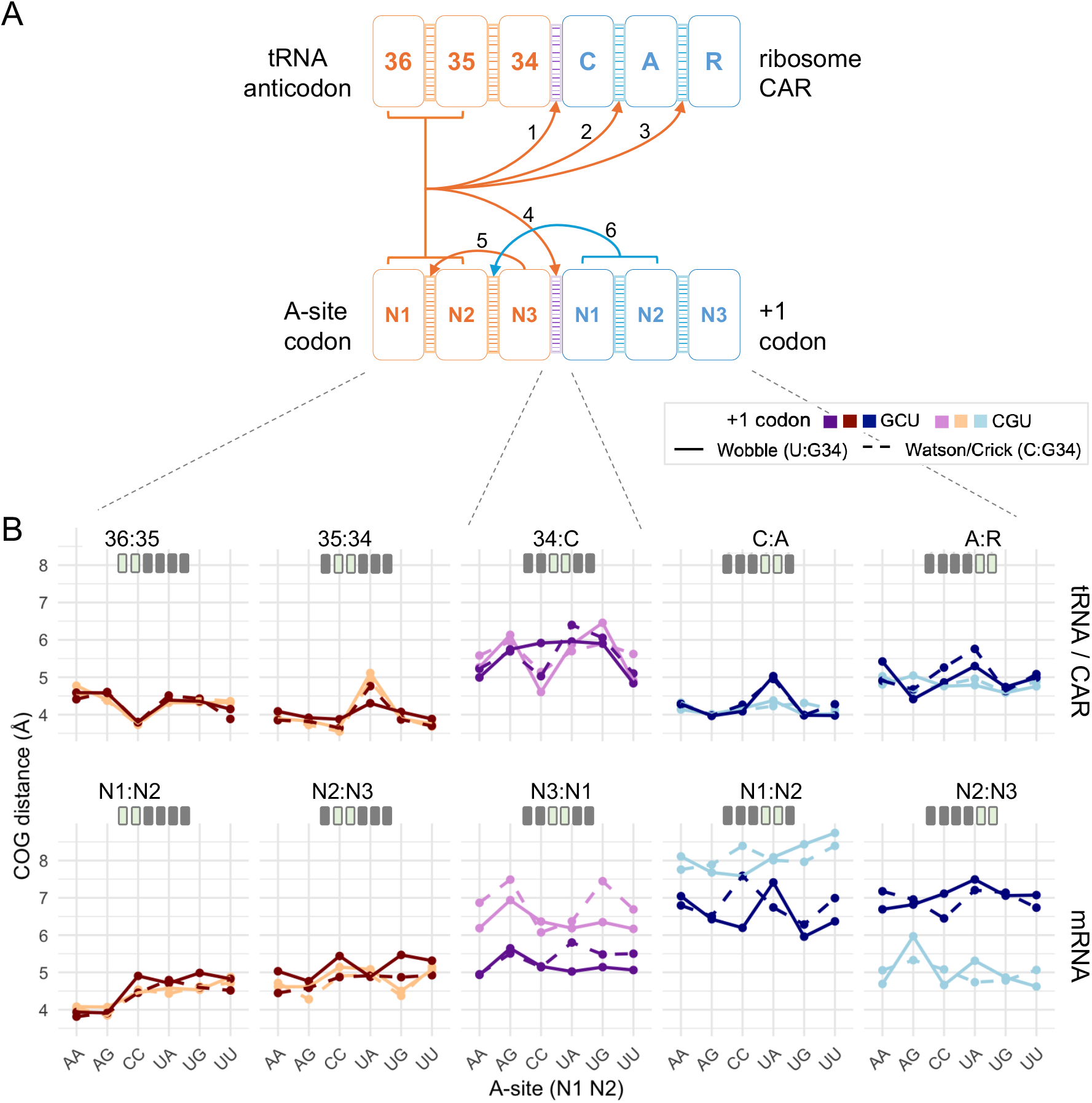
Structural network and codon-dependent variation of stacking interactions across the A site and CAR site. (**A**) Schematic of the 10 adjacent nucleotide stacking interactions spanning the tRNA anticodon, A-site mRNA codon, CAR surface, and +1 codon. Arrows represent all the non-zero-order effects (p < 0.01) that are one or two residues away from the varying nucleotide. From left to right the stacking residue pairs are: 36:35 and 35:34 (internal anticodon contacts), 34:C (anticodon : CAR tether), C:A and A:R (internal CAR contacts) for the top row, and N1:N2 and N2:N3 (internal A-site codon contacts), N3:N1 (inter-codon bridge), N1:N2 and N2:N3 (internal +1 codon contacts) for the bottom mRNA row. All A-site codon – anticodon nucleotides and interactions are in hues of red, inter-site stacks in hues of purple and CAR - +1 codon region nucleotides and interactions are in hues of blue. (**B**) Center-of-geometry distances (Å) measured for the same ten stacking interactions, plotted for each A-site N1N2 codon (x-axis). Dark lines: +1GCU context; light lines: +1CGU context. Solid traces denote G:U wobble pairing at the A-site third position; dashed traces denote G:C Watson–Crick pairing. Color coding matches panel A.

Internal stacking within the tRNA anticodon (36:35, 35:34) and A-site mRNA (N1:N2, N2:N3) showed consistently low COG values (good stacking) (Fig. 3B, Fig. S8). These stacking events reproduce known stacking behaviors and validate the accuracy of our analysis pipeline. Nonetheless, even these canonical A-site codon and anticodon stacks exhibited modest sensitivity to A-site identity (zero order effects), underscoring the tRNA’s ability to register codon identity through fine-grained structural readouts beyond reading base pairing.

Striking non-zero order effects across the neighborhood were observed. Stacking interactions within CAR itself (C:A and A:R) were highly responsive to upstream A-site codon identity (arrows 2 and 3, Fig. 3A); e.g., COG aCCU_+1GCU < aUAU_+1GCU (Fig. 3B), as well as to the +1 codon identity e.g., COG aUAU_+1GCU > aUAU_+1CGU (Fig. 3B). These higher-order effects— where A-site codons such as aUAU show pronounced modulation of internal CAR stacking— suggest that CAR senses A-site and +1 codon nucleotides and reshapes its geometry in response to them. In contrast, stacking of +1 N1:N2 and +1 N2:N3 do not show significant sensitivity to A-site context. These results position CAR as an integrator of mRNA adjacent codon context rather than a passive static scaffold.

Context sensitivity also emerged at the two cross-site stack pairs: 34:C, anchoring the tRNA anticodon to CAR, and N3:N1, bridging the A-site codon and the +1 codon. A-site mediated changes to 34:C (arrow 1, Fig. 3A), hints that the A-site mediated changes to the CAR site may be regulated via anticodon : CAR anchoring. While passive participation of CAR cannot be ruled out, altogether, these observations suggest that the 34:C:A:R stacking is likely the structural conduit essential to the interplay between the A and CAR sites. Moreover, the +1 codon mediated changes at the A-site are likely regulated in part through the modulation of A-site mRNA N2:N3 stacking (arrow 6, Fig. 3A) as observed in COG aUGU_+1GCU > aUGU_+1CGU (Fig. 3B). Thus, the aN2:aN3 interaction may belong to a structural signaling pathway through which the CAR site feeds into the A site. The statistical significance matrix in Table S5 reveals a spatial gradient in the influence of codon identity on stacking. These effects dilute at the distal sites (e.g., +1N1:N2, +1N2:N3). Thus, the interactions surrounding 34:C:A:R and aN2:aN3 seem to be regulating this codon-coupled network.

The decoding center may operate through a form of localized allostery, in which steric or electrostatic perturbations are transmitted through π-stacking continuity across the 34:C:A:R column. The stacking effects observed here may actively modulate the H-bonding patterns of the interplay—either as regulators of the coupling itself or as manifestations of a broader regulatory circuit involving additional residues in the ribosomal network. Such a circuit might communicate through a combination of H-bonding and long-range electrostatic polarization along with stacking, allowing the ribosome to integrate codon context via multiple, interlinked physical channels.

While CAR’s role in the 34:C:A:R stacking—the likely mediator of A-side codon effects at the CAR site—was clearly evident, its role in the CAR site mediated changes to the A site, especially the modulation of aN2:aN3 remained unresolved. We hypothesized that the effect of the +1 codon identity at the A site could also be tuned by CAR rather than independently transmitted via mRNA stacking alone. In this scenario, CAR would be an active regulator of the A site decoding center rather than a passive bystander. To explore this possibility, we isolated individual stacking and H-bonding interactions at both sites. As shown in the figures 4C, 4D, and 4E, these interactions were indexed as labeled interactions from “a” to “z”.

**Figure 4.**
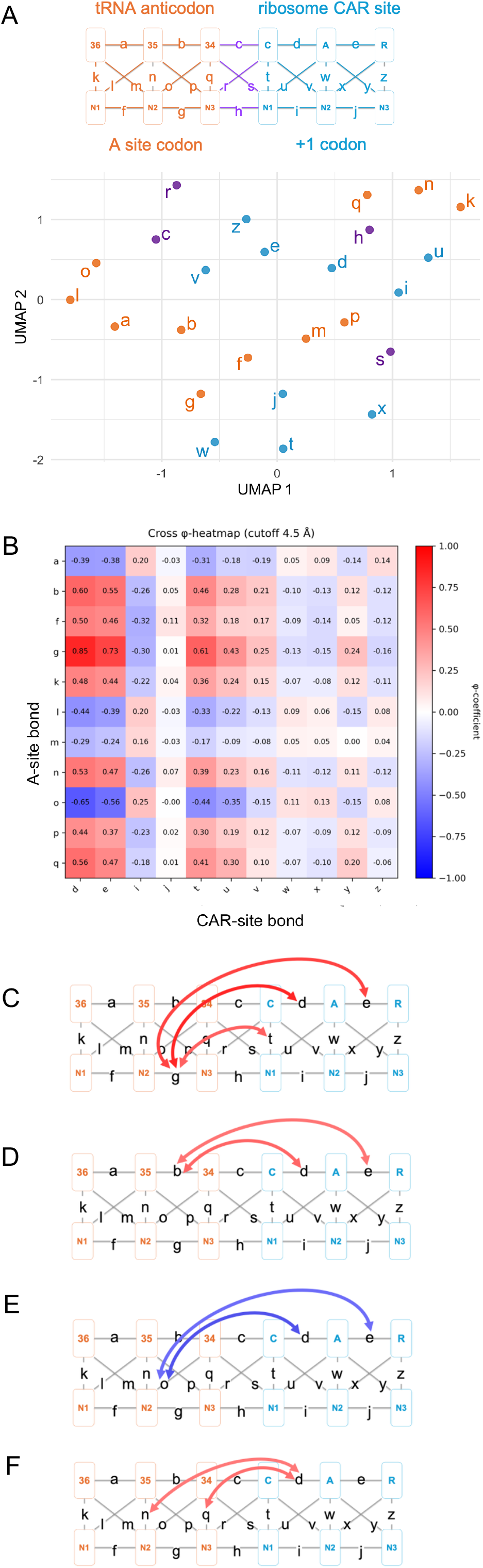
Structural connectivity and covariation across ribosomal A-site and CAR-site regions. (**A**) UMAP clustering of all H-bonding and stacking interactions labeled from “a” to “z”. Each point represents one interaction type embedded in the 2-dimensional UMAP space based on its correlation across different structures. Bond colors indicate site of origin: A-site H-bonds and stacks in orange, CAR and +1 codon interactions in blue, and interface contacts in purple. Each bond’s vector has 720 elements corresponding to 720 simulations (24 structures X 30 replicates). (**B**) Heatmap showing the Phi correlation (ϕ) between A-site and CAR-site interactions. Red and blue shading indicate positive and negative correlation strength. CAR site interactions d and e emerge as cross-site hubs, showing strong correlations with multiple A-site interactions. (**C–F**) Schematics of the decoding center neighborhood with arrows indicating the strong cross-site bond correlations (|ϕ | > 0.5). Arrows are colored according to the correlation’s direction (positive = red and negative = blue) and strength shown in (B).

Each structural configuration (total 24 structures) was clustered using Uniform Manifold Approximation and Projection (UMAP) based on its overall interaction strength profile. This interaction strength profile is a vector including 26 entries corresponding to the quantitative strength of a through z interactions shown in Fig. 4A for that structure (Fig. S9). Distinct clustering patterns emerged based on A-site identity and +1 codon context, consistent with our experimental setup.

The data set in this study provides a behavior perspective of pairwise, residue interactions—H-bonding and stacking—for 24 different versions of the structure with various codons at the A site and CAR site, with 30 MD simulation replicates of each version. This allowed us to create an interaction strength behavioral profile for each pairwise interaction (total 26 interactions labelled as a through z), represented as a vector of 24×30 measurements (720 vector entries for each residue pairwise interaction).

Embedding these vectors into two-dimensional space using UMAP and PCA analysis, based on Pearson correlation measurements between pairwise interaction comparisons of a through z, revealed clustered regionalization of these vectors according to residue interactions within the A site (red; Fig. 4A, Fig. S10), CAR site (blue) or interactions across the A site CAR site boundary (purple). However, the close proximities of vectors at the perimeters of neighboring cluster regions (Fig. 4A, Fig. S10) suggests correlated behaviors of A site interactions with CAR site interactions, even though these interactions are distant in physical space, consistent with our general observations that there are neighborhood-wide correlating behaviors across the ribosome decoding center.

Using frame-resolved binary interaction data, we computed φ correlations between all H-bonding and stacking interactions spanning the A site, CAR site, and +1 codon region (Fig. 4B, Fig. S11). This analysis revealed strong and structured cross-site correlations, suggesting coordinated interaction behavior across the decoding center neighborhood. Several CAR-site interactions emerged as cross-site hubs, exhibiting strong correlations (|φ| > 0.5) with multiple A-site stacking and H-bonding events (Fig. 4B–E). Notably, CAR’s internal stacks (C:A and A:R) showed consistent coupling with A-site interactions, despite physical separation between the residues involved. This underscores CAR’s role in the interplay.

Our results suggest that C of CAR may fine tune the aN2:aN3 stack’s modulation via it’s internal stacking and H-bonding with the mRNA (Fig 4C). Rather than acting as a passive conduit, CAR seems to be a mediator that transmits regulatory signals in both directions by not only integrating upstream input from the A-site but also by relaying +1 codon’s effect back to the A site. The g ⟷CAR correlation—linking A-site stacking (aN2:aN3) to CAR stacking (C:A and A:R) and H-bonding (C:+1N1)— underscores how distinct thermodynamic interactions, such as stacking and H-bonding, could act in concert to enable precise molecular coordination.

Additional cross-site interaction correlations further suggested CAR’s contribution to the interplay (Fig. 4B-F). Bonds b, o, n and q, representing stacking and H-bonding within the A site also showed strong correlations with CAR. These correlations with A-site interactions suggest a regulatory networking where the CAR interaction surface mediates structural effects at the A site. These findings support a model where CAR functions not merely as a local interaction interface but as a regulatory node mediating the interplay between adjacent codons through an integrated network of stacking and H-bonding between the CAR and A sites.

## CONCLUSIONS

By characterizing H-bond and stacking interactions between residues in the ribosome’s A and CAR sites using MD simulations, we have uncovered quantitative relationships that suggest an interplay of signaling between the A site and CAR site that likely contributes importantly to ribosome function. We found an adjacent site interplay wherein the A site affects the CAR site H-bonding and therefore potentially its braking system functionality inferred from analysis of ribosome profiling data(17). Conversely, we also observed that the CAR site affects the A-site H-bonding and potentially codon recognition and frame maintenance. The CAR interaction surface through adjustments of its stacking behavior, appears to be a regulatory node in this interplay, that interprets and relays the information in the structural architecture of the A site, including its wobble geometry, and the +1 codon. Our MD analysis of a Taura syndrome viral mRNA/yeast ribosome subsystem support a model of translation where the ribosome machinery decodes adjacent codon sequences in addition to individual codons. The H-bonding and stacking interplay between the A and CAR sites might serve as a structural basis through which the ribosome manifests the cis-regulatory signals embedded within adjacent mRNA codon sequences. The observations that nt 34 of the tRNA anticodon, which stacks with C of CAR, can be modified under different cellular conditions *in vivo* raises the interesting possibility that the interplay may also be tuned by regulation of these modifications(46–50).

Our findings resonate with a growing body of functional evidence supporting our model that the CAR surface can serve as a critical regulatory interface for tuning the H-bonding base pairing at the A site during translation. Among its residues, C1054 has long been implicated in translation fidelity through genetic studies involving point mutations and suppressor screens(24–28). These studies link C1054 to codon recognition, frame maintenance, and termination at the A site, yet the structural basis for these roles had been unclear. Moreover, the bacterial toxin HigB stacks directly with C1054 to enforce high codon selectivity, in contrast to broader-spectrum toxins like RelE and YoeB, which bypass C1054 and exhibit less specificity(29). The bacterial GTPase RsgA, involved in 16S rRNA maturation, modulates the region surrounding C1054, potentially shielding it and thereby affecting A-site tRNA accommodation(30).

Pharmacological studies further support the functional significance of CAR’s crosstalk with the A site. Antibiotics like tetracycline and tigecycline inhibit decoding through stacking and H-bonding with C1054, and resistance proteins such as TetM and TetO directly displace these drugs by competing for the same site(31, 33, 34). Notably, tigecycline’s enhanced resistance profile has been attributed to stronger stacking interactions with C1054, while streptothricin F extends this targeting to A1196, forming base-pair H-bonds that implicate the broader CAR region in drug binding and regulation(32).

Additional regulatory features of CAR extend to R146, which modulates sequence sensitivity during initiation and codon recognition, a feature exploited by SARS-CoV-2 RPS3RS, a viral regulatory element that co-opts host decoding machinery(23, 35). Consistent with the possible involvement of CAR in sequence selection during translation initiation, preinitiation structures with Met-tRNA in the P site show stacked CAR residues interacting with nucleotide 3 of the A-site codon and nucleotides 1 and 2 of the +1 codon (Fig 1F, Fig. S2)(21).

Intriguingly, independent structural studies also validate that information may flow in both directions across the A-site–CAR interface: A-site tRNA binding induces rearrangements in the 30S subunit that promote mRNA positioning across A1196, indicating that tRNA accommodation may directly influence CAR’s mRNA engagement(51, 52). Together, these findings support our model positioning CAR as a bidirectional sensor and effector of local sequence context—capable of interpreting upstream and downstream signals to shape translation dynamics and therapeutic vulnerability.

Knowledge of the interplay between the A site and CAR site may help elucidate fundamental mechanisms of translational regulation during initiation and termination, as well as codon recognition, frame maintenance, and tRNA accommodation during elongation. Additionally, the stacking and H-bonding geometries identified here provide a structural context for designing antibiotics with better resistance profiles through targeting the A-site–CAR interface, where drug binding specificity and efficacy may be sensitive to interactions with C1054, A1196, or R146 of CAR. Beyond explaining longstanding genetic and pharmacological observations at these residues, these insights offer a foundation for exploring how ribosomal architecture interprets and responds to the compositional logic of mRNA.

## ACKNOWLEDGEMENTS

We thank Joseph Coolon, Scott Holmes, Phil Arevalo and Amy MacQueen for discussions, and Henk Meij for technical assistance with high-performance computing. We acknowledge the use of Generative AI (ChatGPT version 4o) for text copy editing and to enhance code where appropriate.

## AUTHOR CONTRIBUTIONS

Mitsu Raval: Conceptualization, Investigation, Data curation, Formal analysis, Methodology, Validation, Visualization, Writing—original draft, Writing—review and editing. Yancheng Zhou: Data curation, Formal analysis, Investigation, Writing— review and editing. Miki Lynch: Data curation, Investigation. Daniel Krizanc: Investigation, Supervision, Writing—review and editing. Kelly Thayer: Investigation, Methodology, Supervision, Funding acquisition, Writing—review and editing. Michael Weir: Conceptualization, Investigation, Methodology, Visualization, Supervision, Project administration, Funding acquisition, Writing—review and editing.

## SUPPLEMENTARY DATA

Supplementary Data are available at NAR online.

## DATA AVAILABILITY

Restart structure and topology files for MD experiments and code for analysis are available at https://doi.org/10.25438/wes02.30090355.v1 and https://github.com/mitsuraval/A-and-CAR-sites-interplay/tree/main.

