## Supplemental Figures and Tables for "Interplay of the ribosome A and CAR sites"

### SUPPLEMENTARY MATERIALS

### SUPPLEMENTARY FIGURES

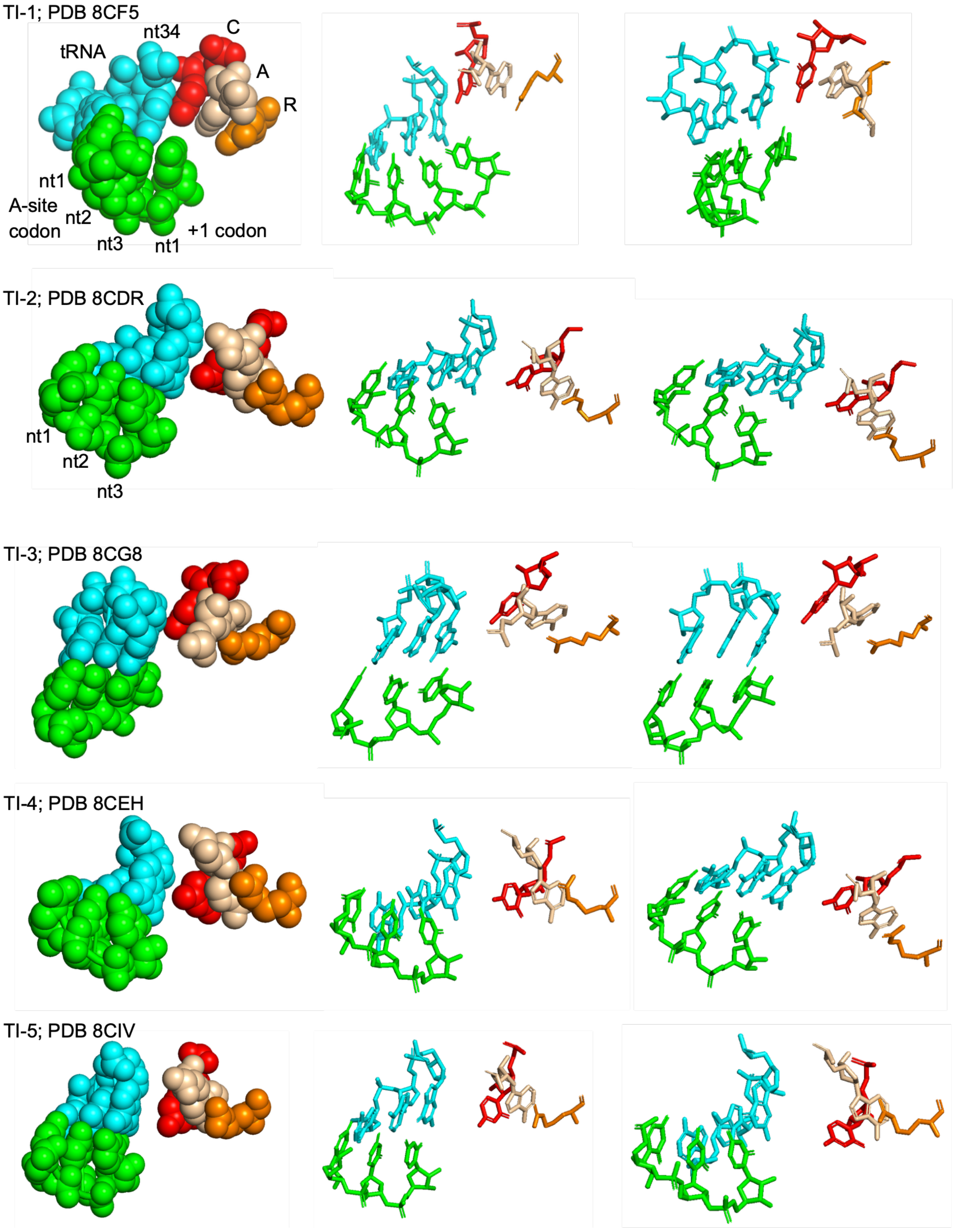


**Figure S1. CAR interface in translocation intermediate structures.**
A- and CAR-site structures in translocation intermediate structures TI-1 through -5 from Milicevic et al. 2024 viewed with sphere (left) and stick (middle and right)

representations. These structures have a tRNA rather than viral IRES. The CAR residues are stacked with each other and in early stages, C (red) of CAR shows stacking with tRNA nucleotide 34 (nt34, cyan). At later stages, CAR is more separated from the tRNA anticodon. TI-1 (PDB ID 8CF5) includes the first nucleotide of the +1 codon 3' adjacent to the A-site codon; since this nucleotide is U rather than G, it is not expected to H-bond strongly with C of CAR. The other translocation intermediate structures (TI-2 through -5) do not include resolved +1 codon nucleotides.

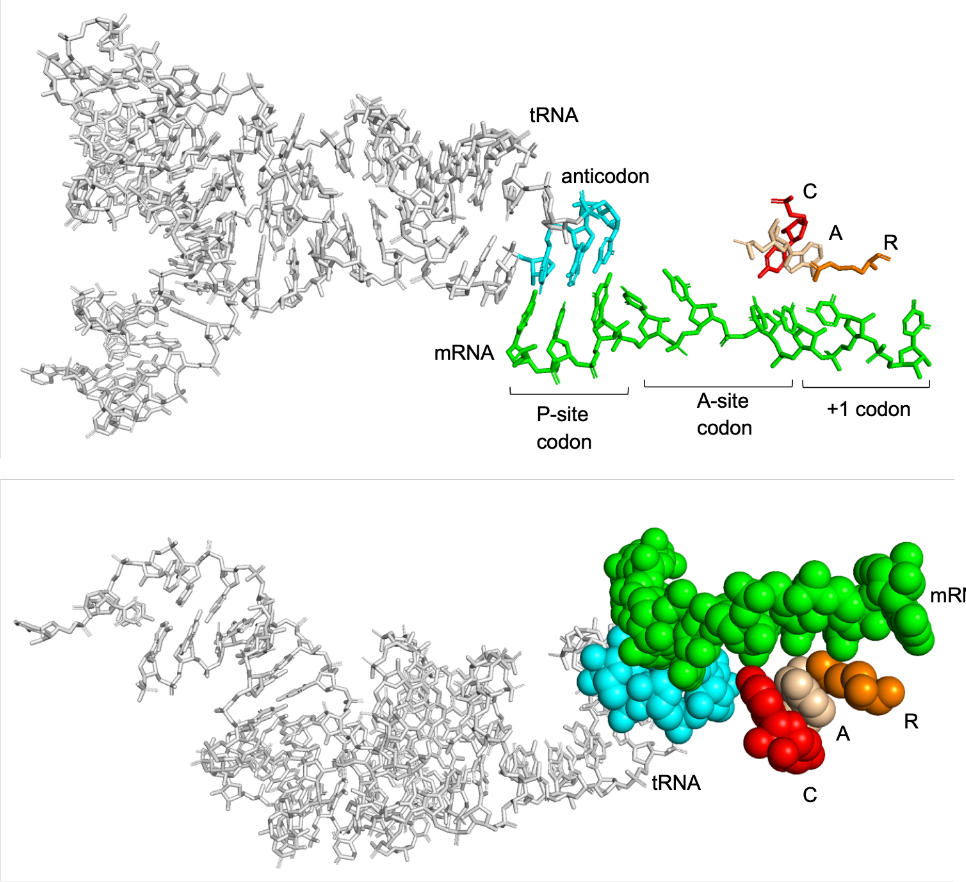


**Figure S2. CAR interacts with mRNA in preinitiation complex.**
Met-tRNA accommodated at the P site in "P-in" preinitiation ribosome (PDB ID 6FYX; Llacer et al. 2018). CAR residues interact with A-site nucleotide 3 (C of CAR), +1 codon nucleotide 1 (A and R of CAR) and nucleotide 2 (R of CAR). The structure is viewed from two angles, showing the stacking of the CAR residues (red, wheat, orange respectively), mRNA nucleotides (green) and tRNA anticodon (cyan).

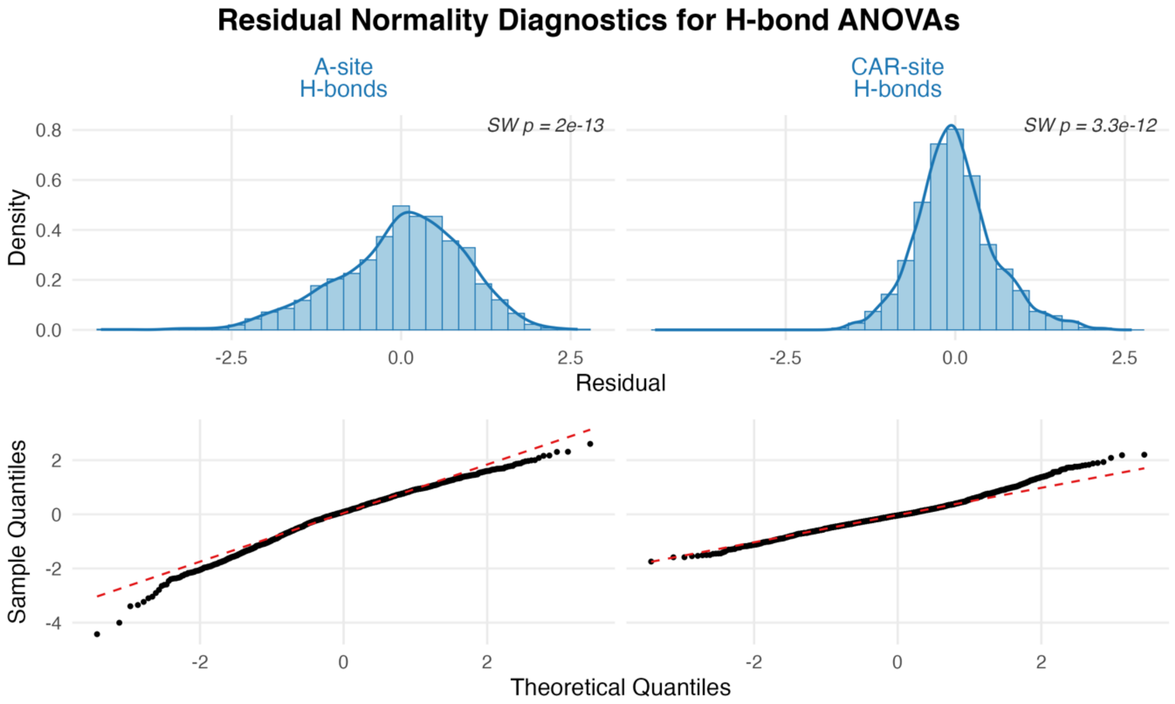


**Figure S3. Residual normality diagnostics for H-bond ANOVAs.**
Residuals from the two four-factor ANOVAs (A-site H-bonds and CAR-site H-bonds) violate the normality assumption, motivating a nonparametric aligned-rank approach. **Top row:** Histograms of residuals (light blue bars with darker-blue border) overlaid with density estimates. Each panel displays the Shapiro–Wilk p-value testing normality—both $p \ll 0.001$, indicating departure from Gaussian distribution. **Bottom row:** Quantile–quantile plots of residuals against a theoretical normal distribution; black points denote observed quantiles, with the dashed red line marking the ideal normal fit. Systematic deviations from the reference line confirm that residuals are not normally distributed in either model.

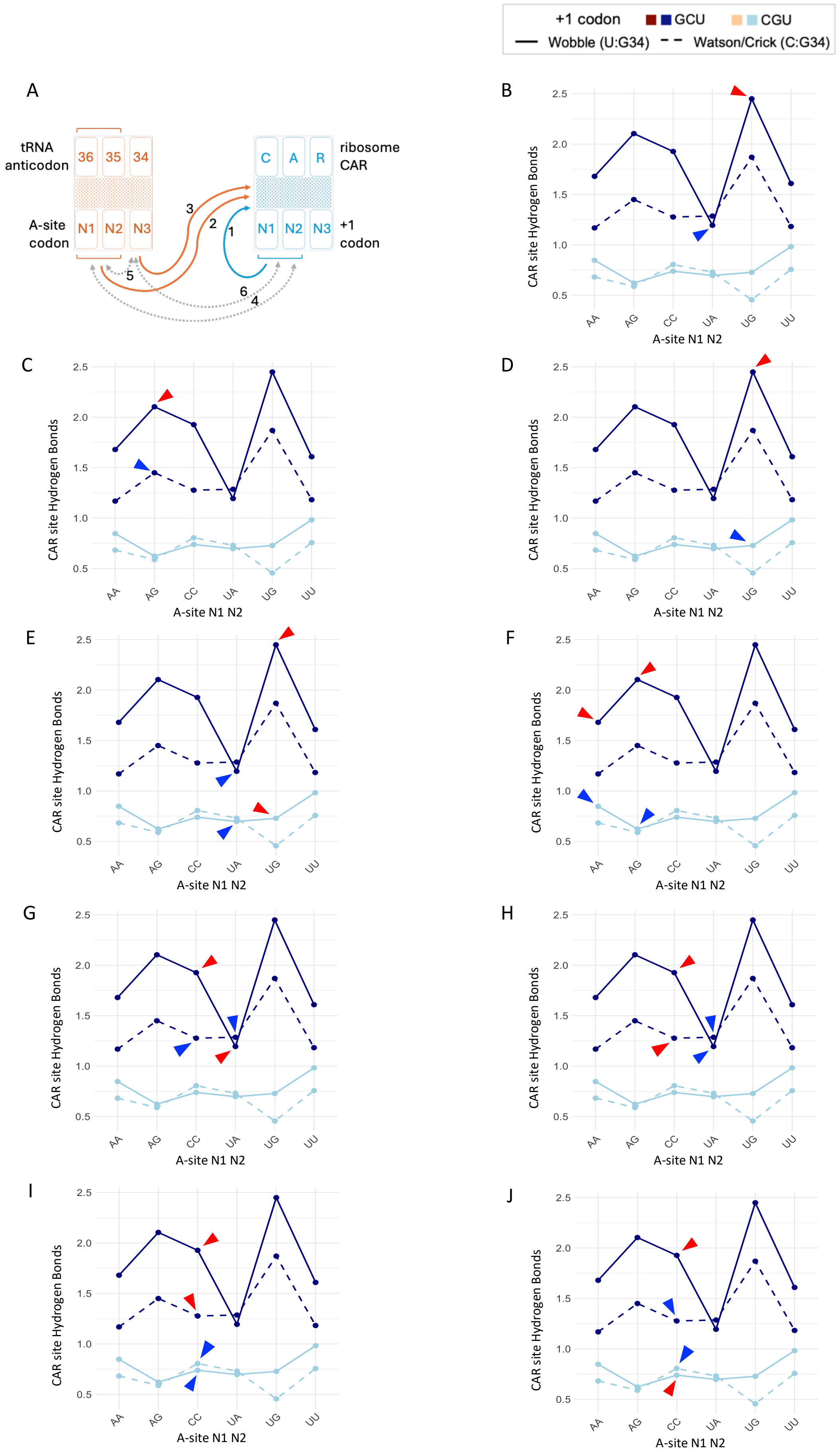

+1 codon GCU CGU
Wobble (U:G34) Watson/Crick (C:G34)
A
tRNA anticodon
36 35 34
A-site codon
N1 N2 N3
ribosome CAR
C A R
+1 codon
N1 N2 N3
1 2 3 4 5 6
B C D E F G H I J
CAR site Hydrogen Bonds
0.5 1.0 1.5 2.0 2.5
AA AG CC UA UG UU
A-site N1 N2

**Figure S4. Replicated panels from Figure 2A and 2B highlighting comparison cases.**

(A) Same as Fig. 2A. (B–J) Each panel shows CAR : +1 codon H-bonding across A-site codons. Blue arrowhead values are always compared relative to red within each subplot to illustrate main and interaction effects.

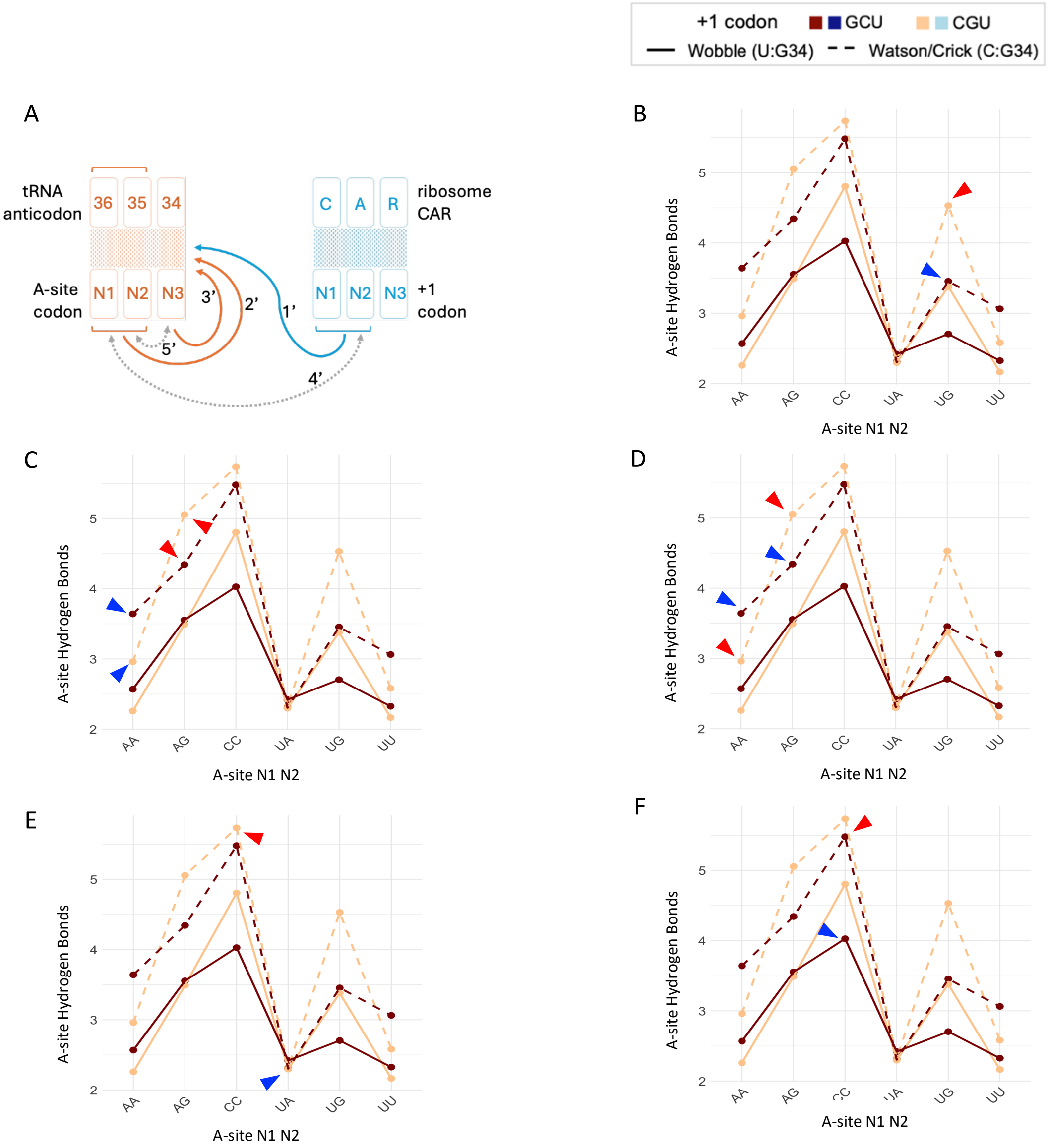


**Figure S5. Replicated panels from Figure 2C and 2D highlighting comparison cases.**

(A) Same as Fig. 2C. (B–F) Each panel shows A-site H-bonding across A-site codons. Blue arrowhead values are always compared relative to red within each subplot to illustrate main and interaction effects.

**A**

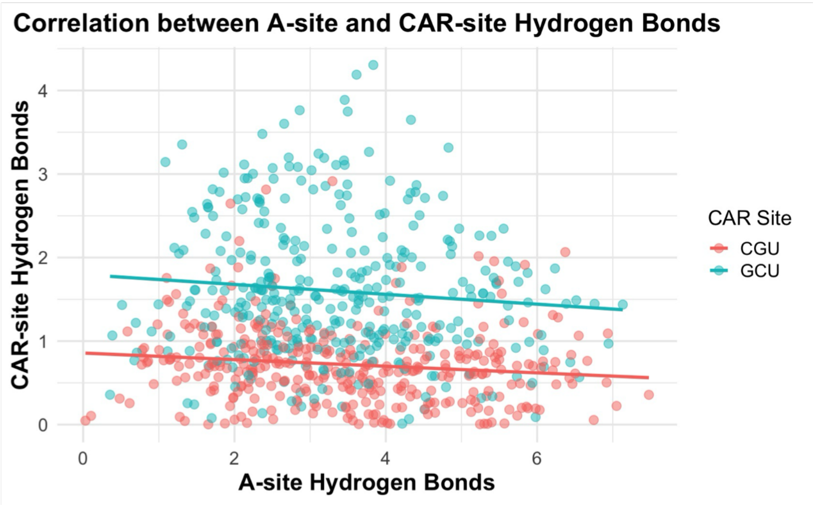


| CAR Site | Pearson r | Raw p-value | Bonferroni p-value | 95% CI |
|---|---|---|---|---|
| CGU | -0.133 | 0.0113 | 0.0227 | -0.2335 to -0.0304 |
| GCU | -0.0956 | 0.0701 | 0.14 | -0.197 to 0.0079 |

**B**

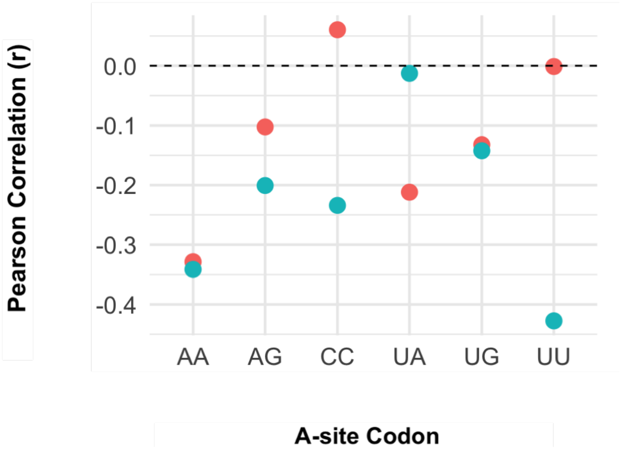


**Figure S6. (A) Correlation between A-site and CAR-site H-bonds.** Each point represents a single molecular dynamics trajectory, showing the average numbers of H-bonds formed between the tRNA and A-site codon (x-axis) and those formed between the ribosome's CAR site and the downstream +1 codon (y-axis). Blue points represent structures with a +1GCU codon, and red points represent +1CGU codons. Lines indicate linear regression fits for each +1 codon group. The observed anticorrelation under GCU conditions suggests that CAR braking is inversely coordinated with A-site decoding strength. Each data point represents results from an MD replicate experiment. **(B) Stratified correlation between A-site and CAR-site H-bonds by A-site codon and +1 codon context.** Each point represents the Pearson correlation coefficient (r) between A-site and CAR-site H-bonding for a specific A-site codon (x-axis), stratified by +1 codon identity (CGU in red, GCU in blue). Negative values indicate anticorrelation between the two sites. The dashed line at r = 0 marks the threshold between positive and negative correlations. Notably, GCU contexts (blue) exhibit stronger negative correlations for certain codons—especially UU—indicating tighter coordination of CAR braking with A-site decoding.

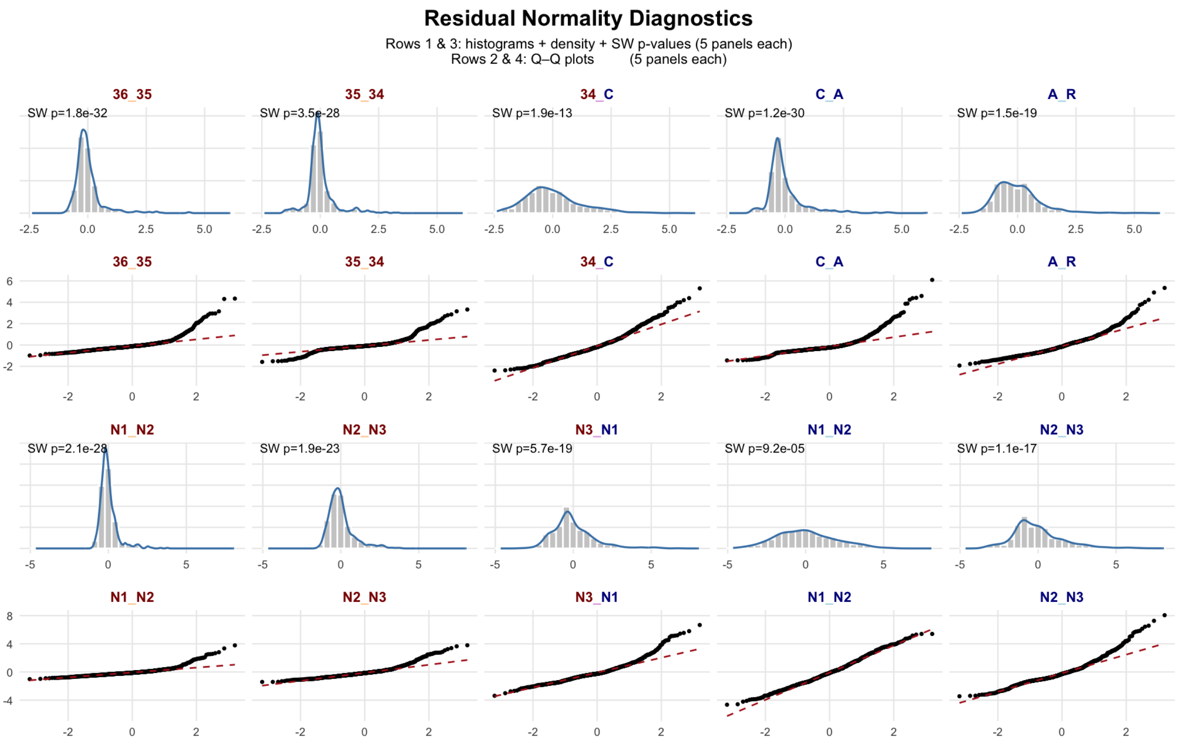


**Figure S7. Residual normality diagnostics for stacking interaction models.** Each column corresponds to a specific stacking interaction assessed in our linear models. Rows 1 and 3 display histograms with overlaid kernel density estimates of model residuals, alongside Shapiro–Wilk test p-values (SW p) for normality. Rows 2 and 4 show corresponding Q–Q plots comparing sample quantiles of residuals to a normal distribution. In all cases, the residuals significantly deviate from normality (SW $p < 0.001$), violating classical ANOVA assumptions. As a result, we employed the aligned rank transform (ART) ANOVA to statistically analyze differences in stacking geometry across A-site and +1 codon contexts.

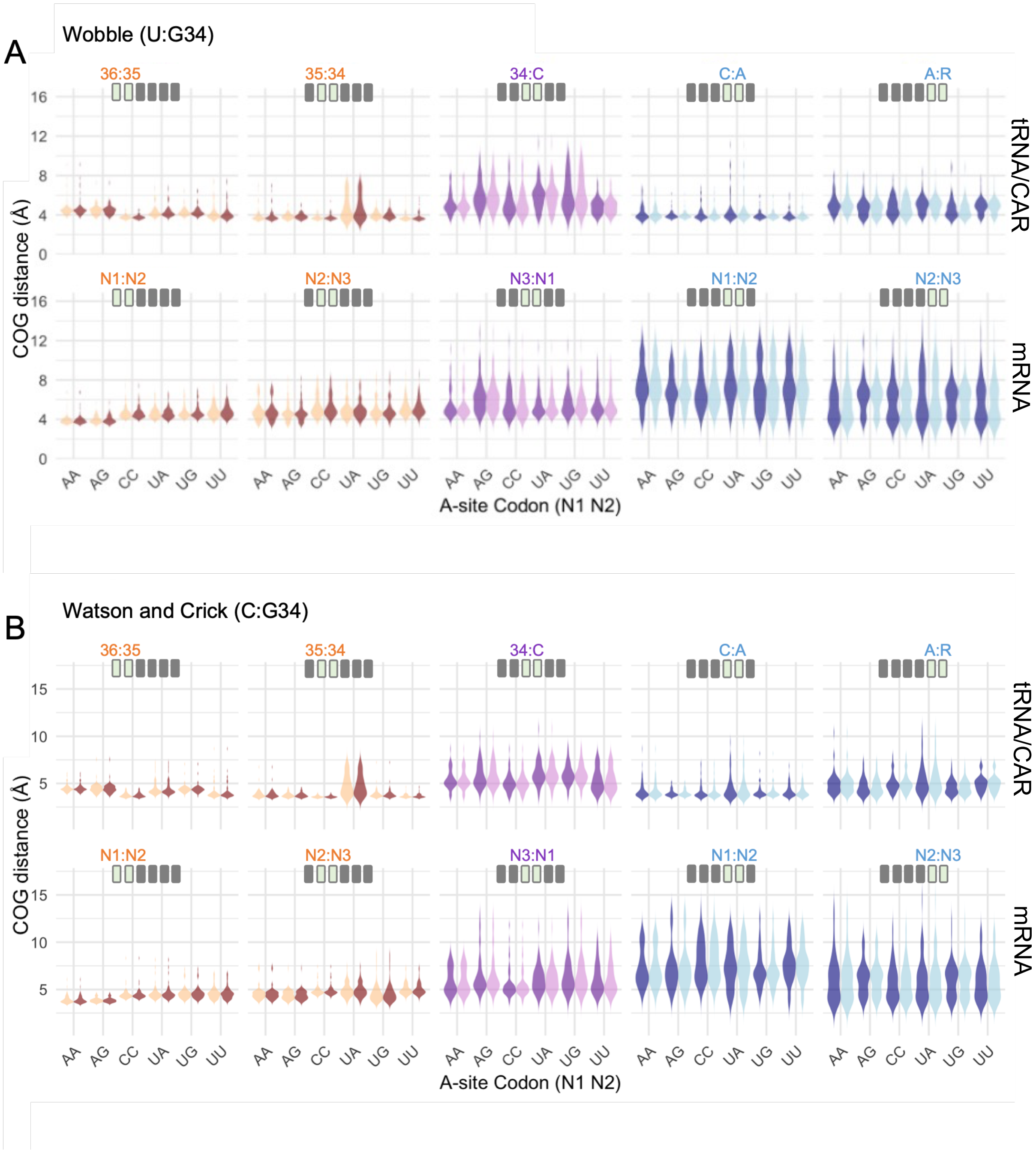


**Figure S8. Stacking interactions across adjacent nucleotide pairs.**
Violin plots depict the distribution of Fig. 3B average center-of-geometry (COG) distances—measured in angstroms—for each nucleotide "stack pair" across the decoding center neighborhood, grouped by the A-site codon (N1N2), for both wobble (A) and Watson–Crick (B) geometries at the third codon position. In each panel, the six A-site codons (AA, AG, CC, UA, UG, UU) are arrayed along the x-axis. The top row of facets shows stacking within the tRNA anticodon (36_35, 35_34), between tRNA anticodon nucleotide 34 and the CAR cytosine (34_C), and internal stacking of the CAR residues (C_A, A_R). The bottom row presents stacking within the A-site codon (N1_N2, N2_N3), the inter-codon bridge from A-site N3 to +1 codon N1 (N3_N1), and

internal stacking of the +1 codon (N1_N2, N2_N3). Color gradients distinguish structural modules: warm tones for tRNA-anticodon stacks, purples for cross-interface interactions, and blues for CAR and +1-codon stacks. Moreover +1GCU and +1CGU correspond to dark and light colors respectively. The width of each violin reflects the full distribution of COG distances observed over 30 molecular dynamics replicates.

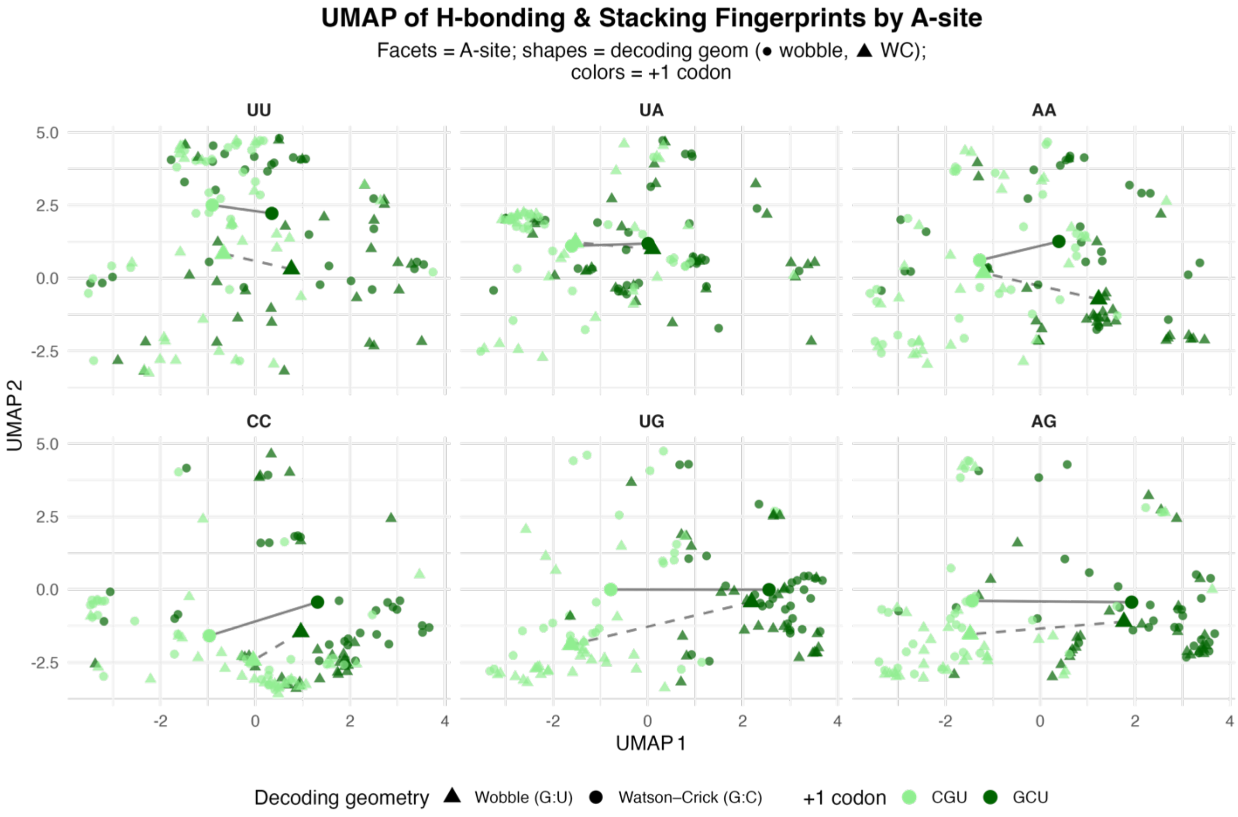


**Figure S9. UMAP of H-bonding and stacking fingerprints by A-site codon.**
Each panel shows all 30 MD simulation replicates for one A-site codon. Points are colored by the +1 codon opposite the CAR surface (dark green = GCU, light green = CGU) and shaped by decoding geometry at the wobble position (● = G:U wobble; ▲ = G:C Watson–Crick). Centroids (larger symbols) indicate the mean UMAP coordinates for each (+1 codon × decoding geometry) group and are connected by gray lines from GCU to CGU within each A-site × geometry facet. UMAP reduces the 26-dimensional hydrogen bonding and stacking fingerprint (bonds a–z) to two axes using a correlation distance matrix (1 – Pearson's r) as input, preserving global covariation structure across frames. Stacking (center-of-geometry) and H-bonding values were scaled using z-scoring prior to distance calculation.

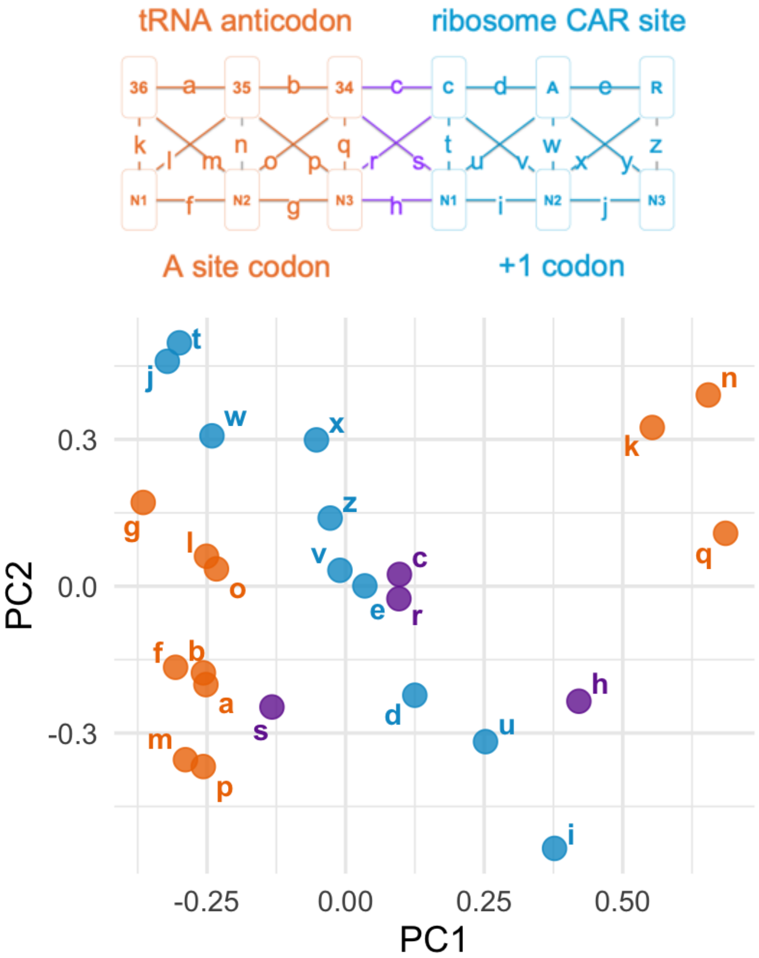


**Figure S10. Principal component analysis (PCA) of structural connectivity across ribosomal A-site and CAR-site regions.**

PCA of all H-bonding and stacking interactions labeled from “a” to “z.” Each point represents one interaction type embedded in the two-dimensional PCA space based on its correlation across different structures. Bond colors indicate site of origin: A-site H-bonds and stacks in orange, CAR and +1 codon interactions in blue, and interface contacts in purple. Each bond’s vector has 720 elements corresponding to 720 simulations (24 structures × 30 replicates).

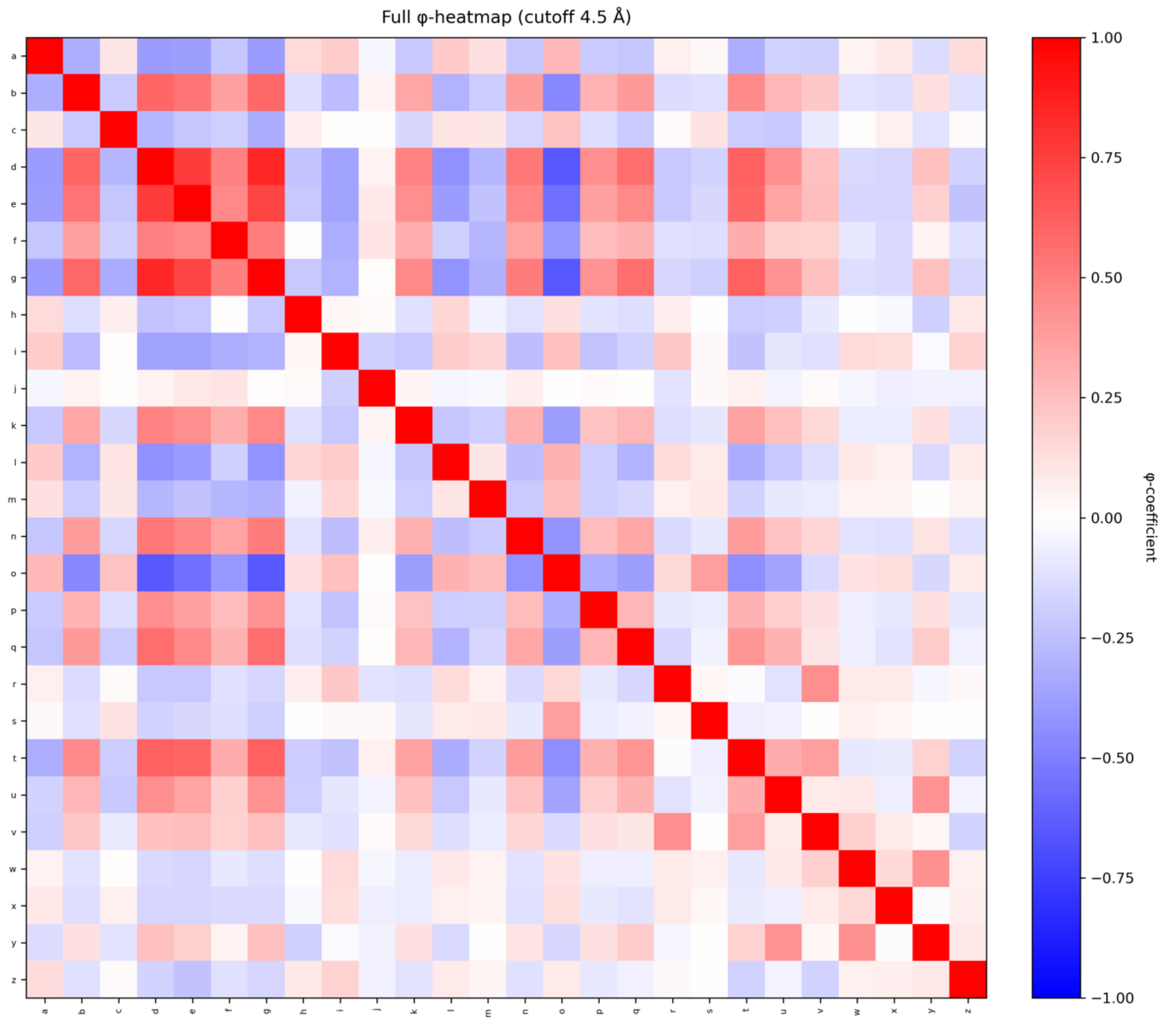


**Figure S11. Full pairwise Pearson-correlation matrix of stacking and H-bonding.** This heatmap displays an extended version of Fig. 4B including all pairwise Pearson correlation coefficients (r) among interaction features (bonds a–z), calculated using mean stacking center-of-geometry distances and H-bonding (based on donor and acceptor atoms distance and angle). The r coefficient captures the degree of covariation between bonds across all MD simulation replicates. Red indicates positive correlation, blue indicates negative correlation, and white represents no correlation. Diagonal elements represent self-correlation (r = 1.0). This matrix reveals site-specific modularity along with intra-site correlations and long-range correlations including the CAR–A-site axis shown in Figure 4.

### SUPPLEMENTARY TABLES

#### Table S1: Summary of ART-ANOVA significance for CAR- and A-site H-bonding[1]

| | A | +1 | W | A↔+1 | +1↔W | W↔A |
|---|---|---|---|---|---|---|
| CAR site | *** | *** | *** | *** | *** | * |
| A site | *** | ** | *** | *** | ns | *** |

#### Table S2: Significant effects of variables on CAR site H-bonding[1]

| | DF | DF.res | Sum Sq | Sum Sq.res | F value | Pr(>F) |
|---|---|---|---|---|---|---|
| +1 | 1 | 696 | 1.210e+07 | 1.889e+07 | 4.458e+02 | 7.464e-76 |
| A | 5 | 696 | 1.397e+06 | 2.956e+07 | 6.579e+00 | 5.397e-06 |
| W | 1 | 696 | 1.727e+06 | 2.924e+07 | 4.112e+01 | 2.641e-10 |
| +1:A | 5 | 696 | 3.489e+06 | 2.745e+07 | 1.769e+01 | 1.582e-16 |
| +1:W | 1 | 696 | 1.070e+06 | 2.985e+07 | 2.496e+01 | 7.422e-07 |
| A:W | 5 | 696 | 4.681e+05 | 3.046e+07 | 2.139e+00 | 5.903e-02 |
| +1:A:W | 5 | 696 | 3.037e+05 | 3.063e+07 | 1.380e+00 | 2.295e-01 |

#### Table S3: Significant effects of variables on A site H-bonding[1]

| | DF | DF.res | Sum Sq | Sum Sq.res | F value | Pr(>F) |
|---|---|---|---|---|---|---|
| +1 | 1 | 696 | 2.8374e+05 | 3.0648e+07 | 6.4435e+00 | 1.135e-02 |
| A | 5 | 696 | 1.4944e+07 | 1.6133e+07 | 1.2894e+02 | 1.356e-96 |
| W | 1 | 696 | 4.6867e+06 | 2.6260e+07 | 1.2422e+02 | 1.169e-26 |
| +1:A | 5 | 696 | 2.1550e+06 | 2.8799e+07 | 1.0416e+01 | 1.185e-09 |
| +1:W | 1 | 696 | 1.3694e+02 | 3.0914e+07 | 3.0830e-03 | 9.557e-01 |
| A:W | 5 | 696 | 1.7235e+06 | 2.9282e+07 | 8.1930e+00 | 1.572e-07 |
| +1:A:W | 5 | 696 | 4.2847e+05 | 3.0483e+07 | 1.9566e+00 | 8.308e-02 |

[1]With reference to Figure 2, these tables S1, S2 and S3 report the statistical significance (aligned rank transform ANOVA) of the three main effects—+1 codon identity (GCU vs. CGU; arrows 1 and 1′), A-site codon identity (N1N2; arrows 2 and 2′), and decoding geometry at the wobble position (G:U vs. G:C; arrows 3 and 3′)—and their pairwise interactions (arrows 4–6 at the CAR site and arrows 4′–6′ at the A site). * p<0.05; ** p<0.01; *** p<0.001. DF = Degrees of freedom. Sum Sq = Sum of squares. Sum Sq.res = Sum of square residuals. These results demonstrate that both CAR- and A-site H-bonding is reciprocally modulated by adjacent codon sequence context.

**Table S4: Covariation of A-site and CAR-site H-bonding by codon context[1]**

| A-site N1 N2 | +1 codon | Pearson's correlation | Raw p values | CI (low) | CI (high) | Bonferroni p values | significant |
|---|---|---|---|---|---|---|---|
| AA | CGU | -0.329 | 0.0104 | -0.538 | -0.0815 | 0.124 | FALSE |
| AA | GCU | -0.341 | 0.00762 | -0.548 | -0.0956 | 0.0914 | FALSE |
| AG | CGU | -0.102 | 0.436 | -0.347 | 0.156 | 1 | FALSE |
| AG | GCU | -0.201 | 0.124 | -0.433 | 0.0561 | 1 | FALSE |
| CC | CGU | 0.0605 | 0.646 | -0.196 | 0.310 | 1 | FALSE |
| CC | GCU | -0.329 | 0.0103 | -0.538 | -0.0820 | 1 | FALSE |
| GA | CGU | -0.212 | 0.104 | -0.442 | 0.0444 | 1 | FALSE |
| GA | GCU | -0.0127 | 0.923 | -0.266 | 0.242 | 0.123 | FALSE |
| UA | CGU | -0.132 | 0.314 | -0.374 | 0.126 | 1 | FALSE |
| UA | GCU | -0.142 | 0.278 | -0.382 | 0.116 | 1 | FALSE |
| UG | CGU | -0.00134 | 0.992 | -0.255 | 0.253 | 1 | FALSE |
| UG | GCU | -0.428 | 0.000656 | -0.615 | -0.195 | 1 | FALSE |
| UU | CGU | -0.329 | 0.0104 | -0.538 | -0.0815 | 1 | FALSE |
| UU | GCU | -0.341 | 0.00762 | -0.548 | -0.0956 | 0.00787 | TRUE |

[1]This table reports Pearson's correlation between the number of A-site and CAR-site H-bonds across all MD replicates for each combination of the first two A-site codon nucleotides (N1 N2) and the +1 codon (CGU or GCU). The table entries list, for each codon pair, the correlation coefficient (r), raw p-value, 95 % confidence interval (lower and upper bounds), Bonferroni-adjusted p-value, and whether the correlation remains significant after correction. Across most codon contexts, no significant correlation was detected after correction. The sole exception is the UUN A-site codons (UU) paired with a +1 GCU codon, which exhibit a negative correlation (r = -0.341, raw p = 0.00762; adjusted p = 0.00787).

**Table S5: Summary of ART-ANOVA significance for stacking (COG distances)[1]**

| Residue pair | A | +1 | W | A↔+1 | +1↔W | W↔A |
|---|---|---|---|---|---|---|
| a36:a35 | *** | ns | ns | ns | ns | ns |
| a35:a34 | *** | ns | ns | ** | ns | ns |
| a34:C | *** | ns | ns | * | ns | ns |
| C:A | *** | ns | ns | * | ns | ns |
| A:R | *** | ns | ns | ns | ns | ns |
| aN1:aN2 | *** | ns | * | ns | ns | ns |
| aN2:aN3 | *** | * | *** | * | ns | ns |
| aN3:+1N1 | ** | *** | ** | ns | ns | ns |
| +1N1:+1N2 | ns | *** | ns | ns | ns | ns |
| +1N2:+1N3 | ns | *** | ns | ns | ns | ns |

[1]This table reports the statistical significance (from Bonferroni corrected p-values) of main effects—A-site codon identity (N1 N2), +1 codon identity (+1GCU vs. +1CGU), and wobble decoding geometry (G:U vs. G:C)—and their two-way interactions on the center-of-geometry (COG) distances for each of the ten nucleotide stack pairs. Stacking interactions at the decoding center and CAR interface are highly context-sensitive and contribute to the bidirectional crosstalk underlying adjacent-codon regulation. * $p<0.05$; ** $p<0.01$; *** $p<0.001$.
